# Cyclin F drives proliferation through SCF-dependent degradation of the retinoblastoma-like tumor suppressor p130/RBL2

**DOI:** 10.1101/2021.04.23.441013

**Authors:** Taylor P. Enrico, Wayne Stallaert, Elizaveta T. Wick, Peter Ngoi, Seth M. Rubin, Nicholas G. Brown, Jeremy E. Purvis, Michael J. Emanuele

## Abstract

Cell cycle gene expression programs fuel proliferation and are dysregulated in many cancers. The retinoblastoma-family proteins, RB, p130/RBL2 and p107/RBL1, coordinately repress cell cycle gene expression, inhibiting proliferation and suppressing tumorigenesis. Ubiquitin-dependent protein degradation is essential to cell cycle control, and numerous proliferative regulators, tumor suppressors, and oncoproteins are ubiquitinated. However, little is known about the role of ubiquitin signaling in controlling RB-family proteins. A systems genetics analysis of several hundred CRISPR/Cas9 loss-of-function screens suggested the potential regulation of the RB-network by cyclin F, a substrate recognition receptor for the SCF family of E3 ligases. We demonstrate that RBL2/p130 is a direct substrate of SCF^cyclin F^. We map a cyclin F regulatory site to a flexible linker in the p130 pocket domain, and show that this site mediates binding, stability, and ubiquitination. Expression of a non-degradable p130 represses cell cycle gene expression and strongly reduces proliferation. These data suggest that SCF^cyclin F^ plays a key role in the CDK-RB network and raises the possibility that aberrant p130 degradation could dysregulate the cell cycle in human cancers.

## Introduction

The eukaryotic cell cycle consists of a sequential progression of events that govern cell growth and division. During cell cycle progression, many hundred genes oscillate in expression, contributing to myriad processes, including DNA replication, chromosome segregation, cytoskeletal organization, etc. The expression of cell cycle genes is highly dynamic but is repressed during quiescence, a reversible state of growth arrest, and in early G1-phase, presenting a critical barrier to proliferation. The retinoblastoma protein (RB) is a vital regulator of cell cycle gene repression. During quiescence and in early G1-phase, RB binds and inhibits E2F transcription factors, repressing transcription of many cell cycle genes. RB is phosphorylated by Cyclin Dependent Kinases 4 and 6 (CDK4/6), as well as CDK2 (Narasimha et al., 2014; Rubin et al., 2020). This phosphorylation causes RB to dissociate from E2F, promoting the E2F-dependent expression of cell cycle genes that catalyze S-phase entry and cell cycle progression (Rubin et al., 2020). Due to its critical role in restricting proliferation, RB is a prototypical tumor suppressor (Dyson, 2016).

RB has two closely related family members, RBL1/p107 and RBL2/p130 (Dyson, 1998; Sadasivam and DeCaprio, 2013). Both p107 and p130 are also tumor suppressors that prevent cell cycle gene expression by binding the repressor E2F proteins E2F4/5 (Claudio et al., 1994; Zhu et al., 1993), and both are also regulated CDK4/6 dependent phosphorylation (Canhoto et al., 2000; Farkas et al., 2002; Hansen et al., 2001). p130 functions as part of the mammalian DREAM complex (<u>D</u>P, <u>R</u>B-like, <u>E</u>2F4/5, <u>a</u>nd <u>M</u>uvB) (Litovchick et al., 2007; Smith et al., 1996).

DREAM assembles during quiescence and inhibits cell cycle progression by restricting the transcription of numerous cell cycle genes regulated by E2F, B-MYB, and FoxM1 transcription factors (Fischer et al., 2016; Müller et al., 2012). Accordingly, perturbations to p130 or the DREAM complex allow expression of its cell cycle target genes, shifting the balance from quiescence towards proliferation (Forristal et al., 2014; Iness et al., 2019; Patel et al., 2019).

RB and p130 collaborate to suppress cell cycle gene expression and proliferation. In mice, *Rb^-/-^p130^-/-^* mouse embryo fibroblasts (MEFs) grow more rapidly in culture than MEFs deficient in either *Rb* or *p130* alone, and *Rb^-/-^p130^-/-^* mice spontaneously form many more tumors than their respective single gene knockouts (Dannenberg et al., 2004). In mouse models of small cell lung cancer, *p130* knockout increases tumor size and overall tumor burden, even in the background of *Rb* and *p53* loss (Ng et al., 2020; Schaffer et al., 2010). Consistent with its role as a tumor suppressor, *p130* cooperates with Rb to repress G2-M genes in response to genotoxic stress (Schade et al., 2019a). And, *p130* loss in primary human fibroblasts leads to increased expression of cell cycle genes compared to loss of *Rb* alone (Schade et al., 2019b). Together, these observations highlight the importance of p130 in cell cycle control, as well as its role in tumor suppression. These results also illustrate the importance of the broader CDK-RB network in normal proliferation, and the consequence of its dysregulation in the aberrant cell cycles observed in cancer. RB mutations, overexpression of cyclin D and cyclin E, loss of p130 protein, and dysregulation of the mammalian DREAM complex have all been implicated in increased cellular proliferation and tumorigenesis (Forristal et al., 2014). Interestingly, p130 mutations are infrequent compared to other tumor suppressors like *RB, CDKNA2*, or *p53*, suggesting the possibility that post-translational mechanisms could account for its inactivation.

Our analysis of large-scale CRISPR/Cas9 loss-of-function screens performed in hundreds of human cell lines suggested a link between cyclin F and the CDK-RB network. Cyclin F is a non-canonical cyclin-it neither binds nor activates CDKs (Bai et al., 1994; D’Angiolella et al., 2013). Instead, cyclin F is one of ∼70 F-box proteins, a family of substrate recognition receptors that recruit substrates to the Skp1-Cul1-Fbox protein (SCF) E3 ligase (Bai et al., 1996; Cardozo and Pagano, 2004). SCF ligases play an evolutionarily conserved role in promoting cell cycle progression by triggering the destruction of cell cycle inhibitors. For example, yeast SCF^Cdc4^ and human SCF^Skp2^ trigger the destruction of CDK inhibitors Sic1 and p27, respectively (Carrano et al., 1999; Feldman et al., 1997; Schwob et al., 1994; Skowyra et al., 1997).

Cyclin F mRNA and protein levels oscillate during the cell cycle, giving cyclin F cell cycle-dependent activity (Bai et al., 1994). Cyclin F begins to accumulate at the G1/S transition, peaks in G2, and its protein levels are subsequently downregulated via proteasomal degradation in mitosis and G1 (Choudhury et al., 2016a; Mavrommati et al., 2018). Cyclin F has been implicated in the ubiquitination and degradation of numerous cell cycle proteins. For example, cyclin F helps keep the Anaphase Promoting Complex/Cyclosome (APC/C) inactive during S-phase by marking its co-activator, Cdh1, for degradation (Choudhury et al., 2016a).

Further, cyclin F is involved in cell cycle gene transcription both through its non-catalytic inhibition of B-MYB after DNA damage (Klein et al., 2015), and its ubiquitination of activator and repressor E2Fs (Burdova et al., 2019; Clijsters et al., 2019; Wasserman et al., 2020; Yuan et al., 2019) and the stem-loop binding protein (SLBP) in G2-phase (Dankert et al., 2016). Together, its dynamic cell cycle regulation and numerous cell cycle substrates highlight the importance of cyclin F in cell cycle control.

In this study, we demonstrate that cyclin F interacts with, ubiquitinates, and regulates the RB-family tumor suppressor p130. We identify the regions in both cyclin F and p130 that are required for their interaction. Interfering with cyclin F-p130 regulation, by using a mutant version of p130 which cannot be ubiquitinated, causes a severe defect in proliferation. Together, these data implicate cyclin F as a new, key player in CDK-RB network regulation, highlighting a critical ubiquitin-based mechanism that controls cell proliferation.

## Results

### Cyclin F/CCNF fitness correlates with the CDK-RB network

To identify genes involved in CDK-RB network regulation, we used a systems genetics approach (Boyle et al., 2018; Doench et al., 2016; Pan et al., 2018; Wang et al., 2017), leveraging the Project Achilles Cancer Dependency Map (DepMap). At the time of our analysis, the DepMap consortium had performed near genome-wide, pooled, CRISPR/Cas9 loss-of function (knockout) screens targeting 17,634 genes in 789 cell lines. Fitness scores, corresponding to each gene in each cell line (17,634 x 789) reflects the relative impact of gene knockout on proliferation and survival. Notably, the loss of genes/proteins in the same pathway or protein complex impacts cellular fitness similarly. Therefore, genes with highly correlated fitness scores are often linked by a genetic or physical interaction (Abramowski et al., 2016; Kory et al., 2020; McDonald et al., 2017; Pan et al., 2018).

The Pearson’s correlation coefficient was computed for the fitness scores between all possible gene pairs (17,634 x 17,634) across all cell lines. Since these screens measure relative growth and survival, they are remarkably useful at detecting interactions and pathways involved in proliferation. For example, among 17,634 genes queried, the gene most highly correlated with the DNA helicase component MCM2 is its complex member MCM4. Further, the two genes most highly correlated with mitogen activated protein kinase MEK1 are its upstream activator BRAF and its downstream effector ERK2. Similarly, genes that encode proteins in the Skp2-Cyclin E/A-CDK2-E2F pathway are highly correlated, as were those involved in autophagy (Fig. S1A). The highly significant correlations between these known interactors validates the general utility of this approach for interrogating signaling networks related to cell cycle and proliferation.

We investigated the CDK-RB network by examining genes whose fitness score correlates with bona fide network components (Fig. S1). The gene most highly correlated with *CDK4* is its coactivator *CCND1/Cyclin D1,* demonstrating that relevant interactors in the CDK-RB network are highly correlated. Interestingly, the 9th most highly correlated gene with *CDK4* was *CCNF*, which encodes cyclin F (Bai et al., 1994; Bassermann et al., 2014). CCNF is also among the top 1% of most highly correlated genes when analyzing to several other key members of the CDK-RB network, including *CDK4*, *CDK6, CCND1, and RBL1* (Fig. S1B).

Accordingly, when we analyze those genes most highly correlated with cyclin F KO, numerous CDK-RB network components score significantly (Fig. 1A). Remarkably, *CDK4, CDK6, CCND1/Cyclin D1, RBL1* and *RBL2* are all among the top 85 genes, out of more than 17,630 (∼0.5%), whose fitness score is most highly correlated with *CCNF*. Thus, the impact of *CCNF* knockout is highly correlated with genes in the CDK-RB network, and similarly the loss of CDK-RB genes is highly correlated with *CCNF*.

**Figure 1.**
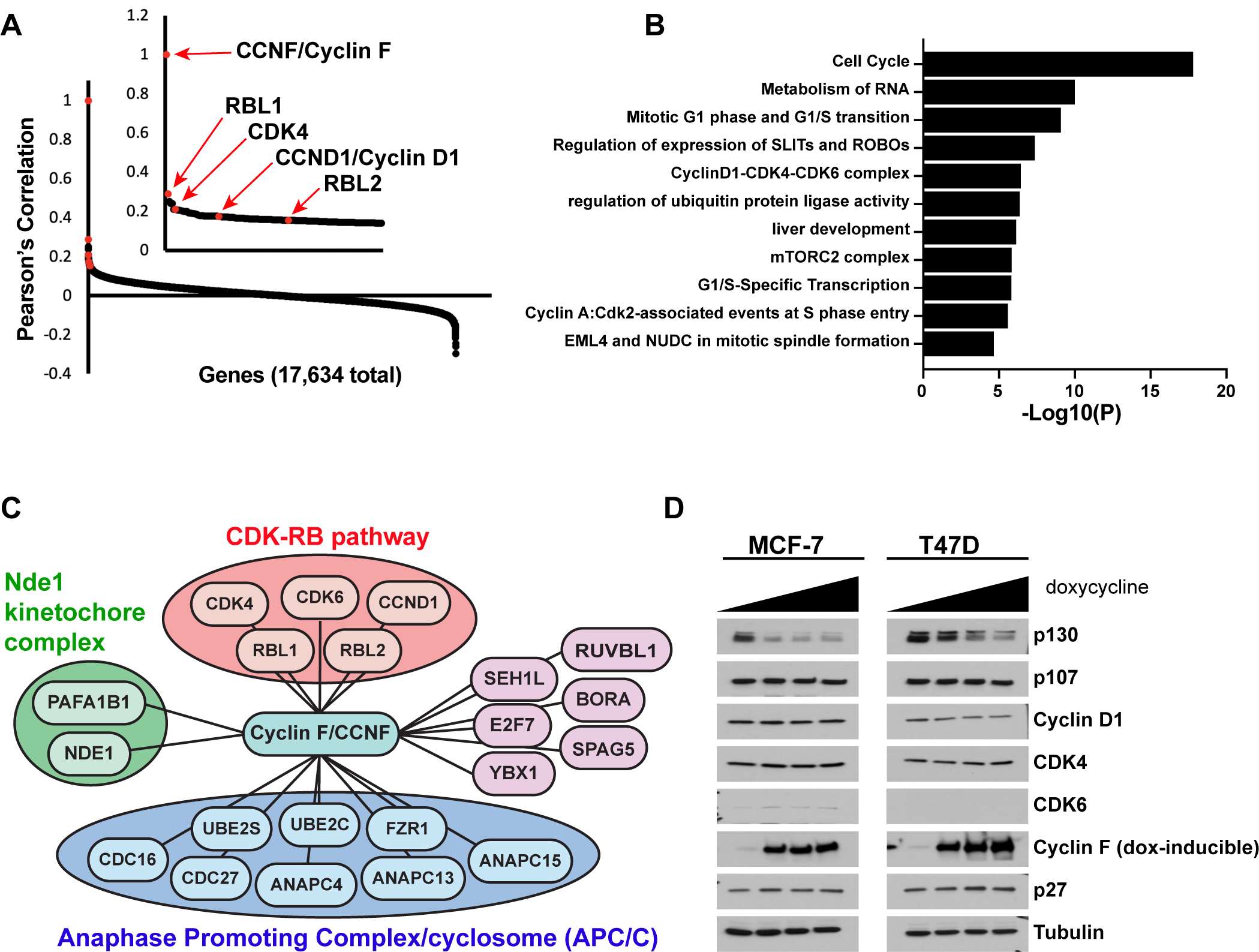
Analysis of the Cancer Dependency Map reveals that *CCNF* is highly correlated with the CDK-RB network. ***(A)*** Cancer Dependency Map data from project Achilles was analyzed to identify impact of gene loss-of-function on cellular fitness, and fitness correlation with that of *CCNF,* based on pooled CRISPR/Cas9 gene knockout screens performed in 789 cell lines. Pearson’s correlation coefficients are reported for all gene pairs (each dot corresponds to a single *CCNF*-gene X pair). The Pearson’s correlations for *CCNF* compared to 17,634 other genes are shown. The CDK-RB network members highlighted in red all score in the top 0.5% of genes whose impact on fitness is most highly correlated with *CCNF*. ***(B)*** Gene ontology (GO) analysis was performed for the top 0.05% of genes whose impact on fitness is most highly correlated with *CCNF*. The top 10 enriched GO terms and their corresponding p-value is shown. ***(C)*** The top 0.05% of genes whose impact on fitness is most highly correlated with *CCNF* were sorted by the GO term cell division (GO:0051301). A graphical representation of the remaining 21 genes is shown. Genes are grouped by their known associations with specific functional pathways or complex, including CDK-RB, Nde1-kinetochore, or APC/C. ***(D)*** MCF-7 and T47D cells were engineered to contain a TET-inducible cyclin F transgene. Cells were treated with increasing amounts of doxycycline to induce cyclin F expression, and the indicated proteins were analyzed by immunoblot. (representative of n=3 experiments)

We performed gene ontology analysis on the top 94 genes whose fitness scores were most strongly correlated with *CCNF* (Pearson correlation >0.15) (Fig. 1B). The GO terms cell cycle and regulation of ubiquitin protein ligase activity are both highly significant. Also enriched are GO terms related to proliferation, including mitotic G1 phase and G1/S transition, Cyclin D1-CDK4-CDK6 complex, G1/S specific transcription, and Cyclin A-CDK2 associated events at S-phase entry. Cyclin F has previously been linked to the cell cycle, as its mRNA and protein levels are cell cycle regulated (Bai et al., 1994), and it is known to mark cell cycle proteins for degradation. However, cyclin F has not previously been linked to regulation of the CDK-RB network.

When we filtered the top *CCNF* correlated genes by the GO term ’cell division’ (Fig. 1C, S1C), three sub-networks emerged: the Nde1-kinetochore complex, the Anaphase Promoting Complex/Cyclosome (APC/C), and the CDK-RB network. Genes encoding two known cyclin F substrates also emerged: *E2F7*, which encodes the transcriptional repressor E2F7 (Yuan et al., 2019), and *FZR1*, which encodes the APC/C coactivator protein, Cdh1 (Choudhury et al., 2016b). Significantly, identification of eight subunits of the APC/C complex is consistent with our previous observation of a reciprocal relationship between SCF^cyclin F^ and APC/C^Cdh1^ (Choudhury et al., 2016a).

We next examined endogenous proteins corresponding to the CDK-RB network genes that correlate most strongly with *CCNF*. We used two breast cancer cell lines, MCF-7 and T47D, engineered for doxycycline-inducible expression of cyclin F (Wasserman et al., 2020). Cells were treated with increasing amounts of doxycycline for 24 hours, then harvested for immunoblot to determine levels of CDK4, CDK6, cyclin D1, p107, and p130. We observed a specific downregulation in p130 protein levels as cyclin F levels increase in both cell lines, whereas all other proteins remained unchanged (Fig. 1D). This downregulation suggests that cyclin F might regulate the DREAM component and transcriptional repressor p130.

### Cyclin F regulates p130

To further assess the relationship between cyclin F and p130, we compared quiescence-synchronized normal human fibroblasts to asynchronous cells. To synchronize in quiescence, cells were serum-starved for 48 hours, then collected for immunoblot analysis. In each of the cell lines tested (BJ, IMR-90, and NHF-1), cyclin F protein levels are high in proliferating cells and low in quiescence, whereas p130 protein levels increase in quiescence, establishing an inverse expression pattern for these proteins (Fig. 2A).

**Figure 2.**
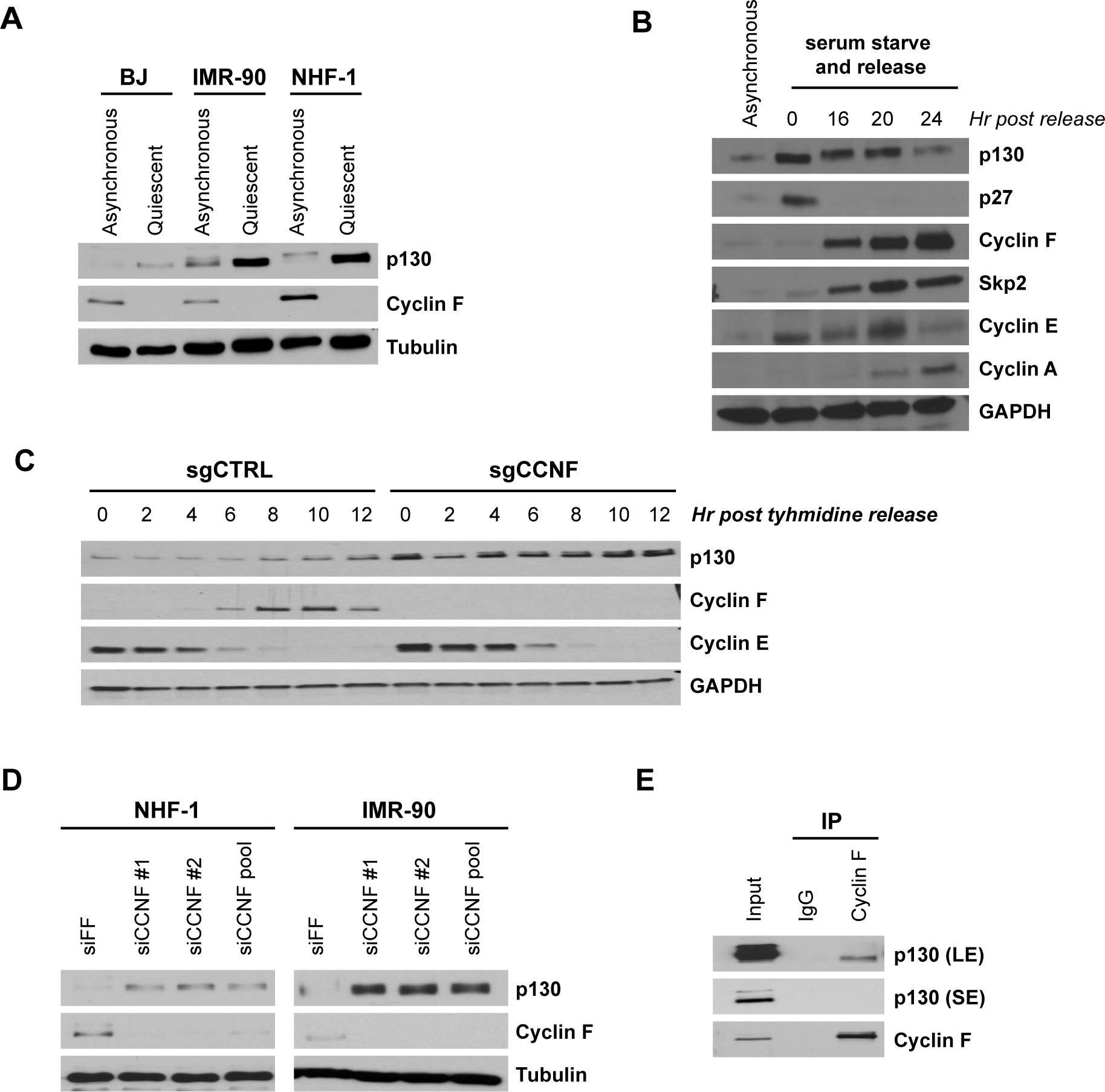
Cyclin F regulates and interacts with endogenous p130. ***(A)*** BJ, IMR-90, and NHF-1 human fibroblast cell lines were synchronized in G0 by 48 h serum starvation (quiescent) or allowed to proliferate normally (asynchronous). Whole cell extracts were collected for immunoblot analysis. No band was detected for CDK6 in T47D cells. (representative of n=3 experiments) ***(B)*** NHF-1 cells were synchronized in G0 by 48 h serum starvation (time 0h) and then released into the cell cycle by re-introduction of serum-containing media. Cells were collected at the indicated times and immunoblotted. Untreated, asynchronous cells (lane 1) are shown as a control. (representative of n=3 experiments) ***(C)*** *CCNF* CRISPR/Cas9 knockouts (sgCCNF) and control (sgCtrl) HeLa cells were synchronized in S-phase by double thymidine block and then released synchronously into the cell cycle upon the addition of drug-free media. Protein levels were monitored at the indicated time points by immunoblot. (representative of n=3 experiments) ***(D)*** NHF-1 and IMR-90 cells were transfected with two different siRNAs targeting *CCNF* or a control siRNA targeting Firefly Luciferase (siFF). Whole cell lysates were immunoblotted for the indicated proteins. (representative of n=3 experiments) ***(E)*** Endogenous cyclin F was immunoprecipitated from asynchronously proliferating 293T cells. Indicated proteins were immunoblotted. (SE=short exposure. LE=long exposure; representative of n=3 experiments).

To determine whether the SCF and cullin E3 ligase family is required for quiescence exit and p130 degradation, we synchronized NHF-1 and T98G cells in quiescence by serum starvation and then released cells into the cell cycle with FBS-containing media supplemented with either DMSO (control) or MLN4924, an inhibitor of the Nedd8 activating enzyme. MLN4924 indirectly inhibits cullins, which require neddylation to be activated (Ohh et al., 2002; Soucy et al., 2009). In cells treated with MLN4924, neither p130 nor p27 are degraded, cyclin E accumulates but does not get degraded, and there is a significant delay in cyclin A accumulation (Fig. S2A). This suggests that cullin-ligases are important for quiescence exit and S-phase entry, since following their inhibition, cells are unable to downregulate p27, which inhibits CDK activation, and p130, which inhibits expression of cell cycle genes through the DREAM complex.

To assess cyclin F and p130 protein levels as cells progress through G1/S, NHF-1 cells were serum-starved for 48 hours to synchronize in quiescence, then stimulated with media containing FBS to allow re-entry into the cell cycle. There was no detectable cyclin F protein in serum-starved cells, but cyclin F protein began to accumulate as cells re-entered the cell cycle at G1/S, similar to what was previously reported (Choudhury et al., 2017; D’Angiolella et al., 2012) (Fig. 2B). Meanwhile, p130 was degraded as cells exited quiescence. The accumulation kinetics of cyclin F is similar to another F-box protein, Skp2, which is responsible for ubiquitinating p27, marking it for degradation. Skp2 has also been implicated in p130 degradation (Bhattacharya et al., 2003; Tedesco et al., 2002). However, p130 has a very different degradation pattern than p27. Moreover, in a previous study, p130 was still degraded in Skp2 knockout cells (Tedesco et al., 2002). Based on the genetic association between cyclin F and the CDK-RB network, the inverse correlation between cyclin F and p130 protein levels, and the ability of cyclin F overexpression to specifically trigger p130 downregulation, we reasoned that SCF^cyclin F^ might control the degradation of p130.

To determine p130 levels in the absence of cyclin F, we first utilized *CCNF* knockout HeLa cells. CCNF KO and control HeLa cells were synchronized in S-phase by double thymidine block. After release, p130 levels were higher in the knockout cells compared to controls at every timepoint analyzed (Fig. 2C). To further investigate the effect of cyclin F loss on p130 levels, we transiently depleted cyclin F in NHF-1 or IMR-90 cells using two different siRNAs. Cyclin F depletion increased p130 protein levels in both cell lines and with both siRNAs (Fig. 2D). Together, these data suggest that cyclin F regulates p130. Consistent with this regulation resulting from a physical interaction, endogenous cyclin F co-immunoprecipitated endogenous p130 from HEK293T cells (Fig. 2E).

### p130 degradation is proteasome- and neddylation-dependent and requires the canonical cyclin F substrate-binding site

We examined if exogenously expressed cyclin F affects the stability of exogenously expressed p130 protein. In HEK293T cells transfected with HA-tagged p130 (HA-p130), increasing amounts of FLAG-tagged cyclin F (FLAG-cyclin F) reduced p130 levels in a dose-dependent manner (Fig. 3A). Four human SCF E3 ligases are involved in marking substrates for degradation to promote entry and progression through S-phase: SCF^cyclin F^, SCF^Skp2^, SCF^βTRCP1/2^ and SCF^Fbxw7^. We therefore also tested whether these other F-box proteins might be able to downregulate p130 levels. We transiently expressed HA-p130 alone or together with cyclin F, Skp2, Fbxw7, βTRCP1 or βTRCP2, and we observed that p130 levels decreased only upon co-overexpression with cyclin F and not with any of the other F-box proteins (Fig. 3B).

**Figure 3.**
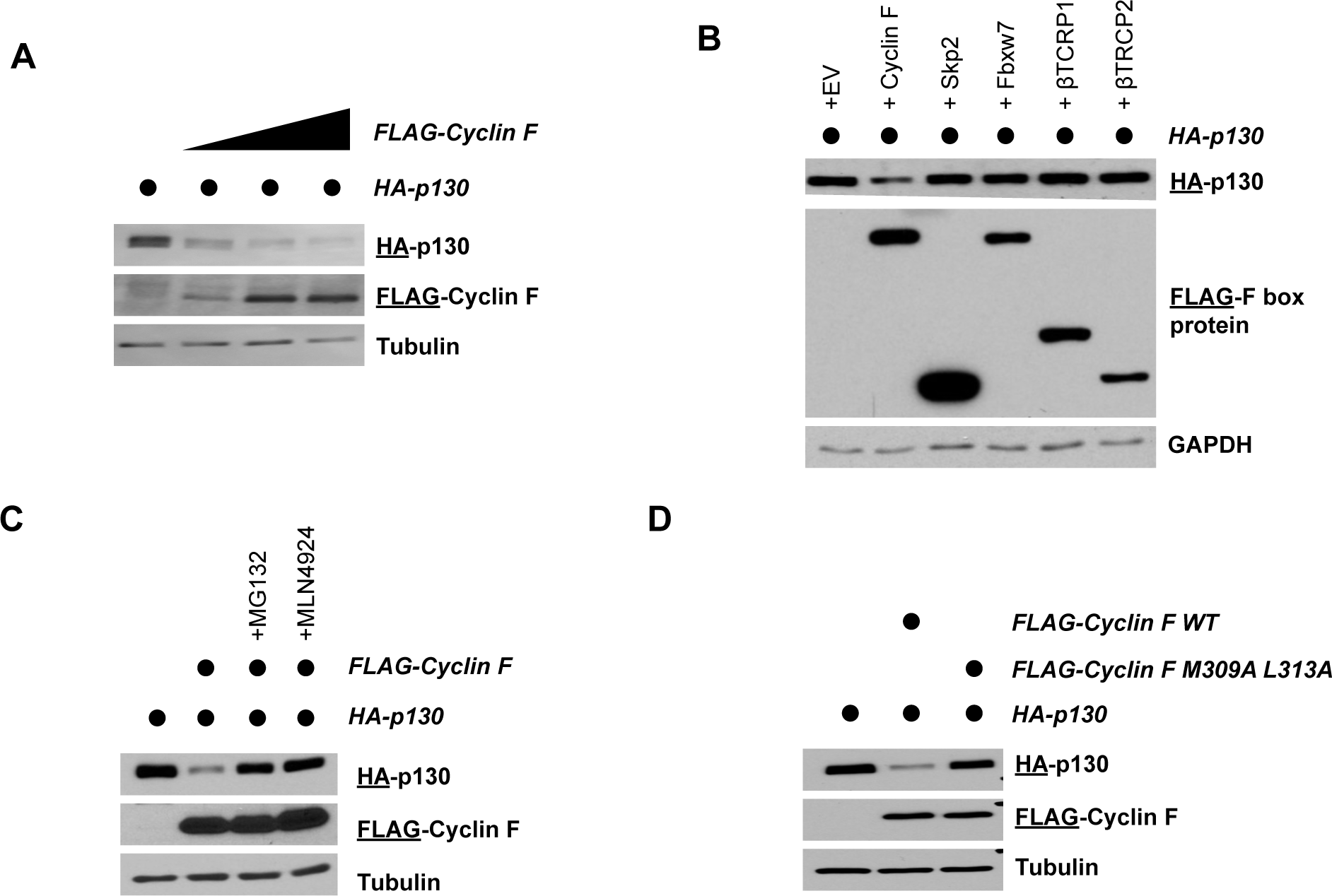
Cyclin F promotes p130 degradation. ***(A)*** HEK293T cells transiently expressing HA-p130 alone (lane 1) or together with increasing amounts of FLAG-cyclin F (lanes 2-4). Cells were collected and analyzed by immunoblot 24h post-transfection. The antigen being immunoblotted for is represented by the underline, here and in all experiments below. (representative of n=3 experiments) ***(B)*** HEK293T cells transiently expressing HA-p130 alone (lane 1) or together with FLAG-cyclin F (lanes 2-4). MG132 (proteasome inhibitor) or MLN4924 (neddylation inhibitor) were added for six hours prior to harvesting. Cells were collected and analyzed by immunoblot 24h post-transfection. (representative of n=3 experiments) ***(C)*** HEK293T cells transiently expressing HA-p130 alone (lane 1) or together with FLAG-cyclin F WT (lane 2) or FLAG-cyclin F(M309A L313A) (lane 3), as indicated. Cells were collected and analyzed by immunoblot 24h post-transfection. (representative of n=3 experiments) ***(D)*** HEK293T cells transiently expressing HA-p130 alone (lane 1) or together with the indicated FLAG-tagged F-box proteins (lanes 2-6). Cells were collected and analyzed by immunoblot 24h post-transfection. (representative of n=3 experiments)

Subsequently, we asked if the observed decrease in p130 protein levels following exogenous cyclin F expression was dependent on the proteasome and neddylation. In HEK293T cells, FLAG-cyclin F expression significantly reduced exogenously expressed HA-p130, and this could be reversed by addition of the proteasome inhibitor MG132 or the neddylation inhibitor MLN4924 for the last six hours before harvesting cells (Fig. 3C). Since p130 degradation is both proteasome and cullin dependent, we determined whether the degradation is also dependent on cyclin F substrate binding. Similar to how canonical cyclins recognize substrates, cyclin F is known to recognize its substrates through a hydrophobic patch motif in its cyclin domain (sequence MRYIL at amino acids M309-L313). Previous studies have shown that mutating the methionine and leucine residues to alanine (Cyclin F M309A L313A) can impair cyclin F binding to substrates (Choudhury et al., 2016b; D’angiolella et al., 2012; D’Angiolella et al., 2010). We transiently expressed HA-p130 with either FLAG-cyclin F(WT) or FLAG-cyclin F(M309A L313A) (Fig. 3D). While cyclin F(WT) robustly downregulated p130 protein levels, cyclin F(M309A L313A) was unable to promote p130 degradation, demonstrating that the previously characterized cyclin F substrate binding, hydrophobic patch motif must be intact to promote p130 degradation in cells. Together, these data suggest that p130 is a cyclin F substrate.

### p130 is a Cyclin F substrate

We next sought to determine binding requirements between p130 and cyclin F. Cyclin F recognizes Cy Motifs in substrates, similar to other cyclins, which correspond to the amino acid sequence RxL/I (where x=any amino acid)(Choudhury et al., 2016b; D’Angiolella et al., 2010, 2012). There are thirteen putative Cy motif sequences in p130 that span the entire length of the protein. We therefore created six HA-tagged p130 truncations, while considering its domain structure (Fig. 4A). We transiently expressed each HA-p130 truncation alone, or together with FLAG-cyclin F (WT). We included conditions with and without proteasome and neddylation inhibitors to confirm that reductions in p130 were cullin ligase and proteasome dependent. We found that p130 degradation depended on the presence of a spacer region, located between sections A and B in the p130 pocket domain, indicating a requirement for this flexible region in cyclin F mediated degradation (Fig. 4A and Fig S3). Within the spacer, there are two RxL/I motifs: R658-I660 and R680-L682. We used site-directed mutagenesis to change the first and last amino acid in each motif to alanine (AxA) in full length HA-p130. We then co-expressed either p130(WT) or the two AxA mutants with FLAG-cyclin F. Only the p130(R680A L682A) mutant was resistant to cyclin F mediated degradation, demonstrating that these two amino acids alone are required for cyclin F to trigger its degradation (Fig. 4B). Notably, that sequence and many surrounding residues are conserved in p130 proteins found in other species, including clawed frogs (Xenopus tropicalis), chickens (Gallus gallus) and mice (Mus musculus) (Fig 4B).

**Figure 4.**
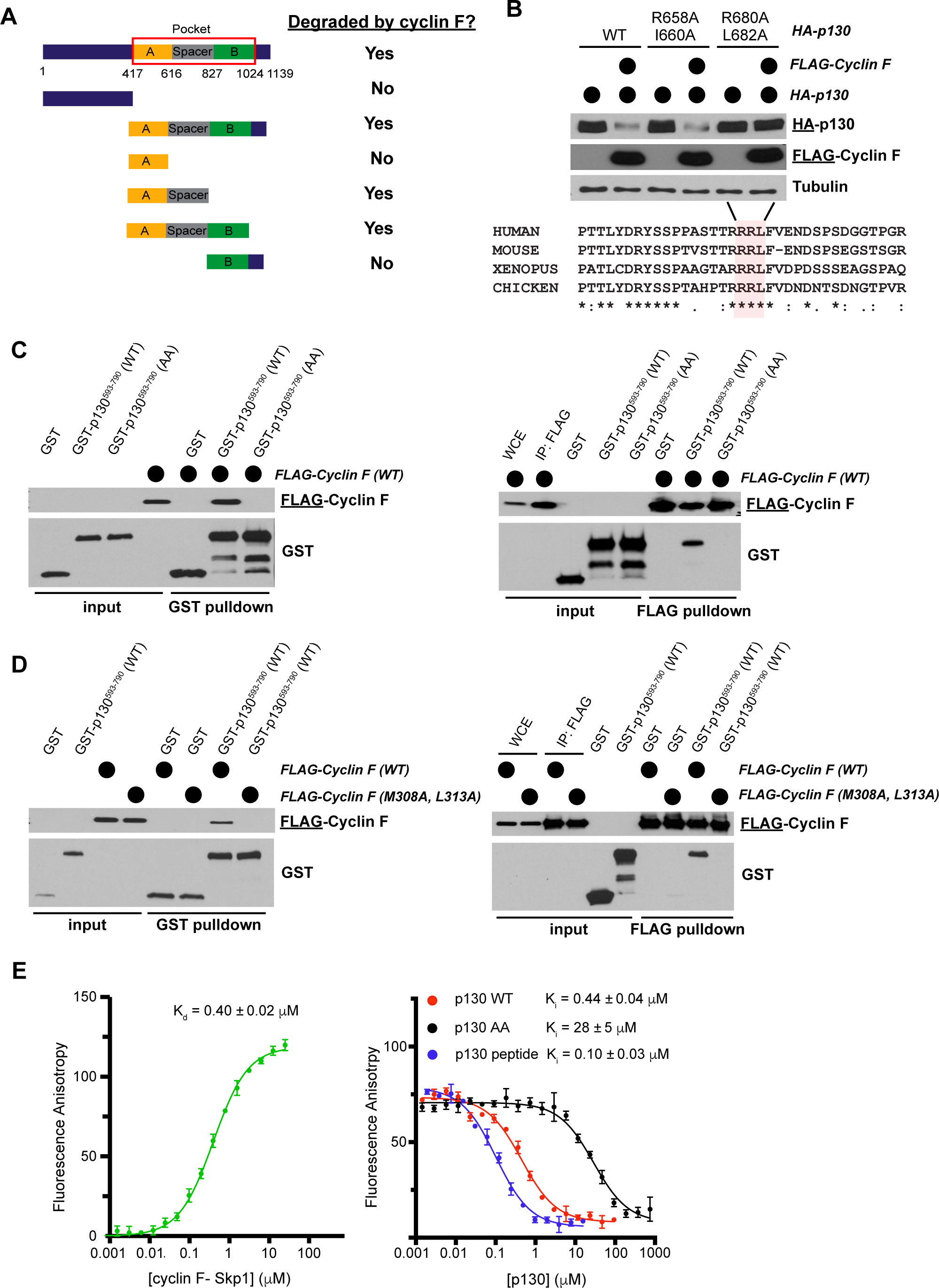
Cyclin F binds p130 directly and promotes p130 degradation through a conserved degron motif. ***(A)*** Graphical depiction of p130 domain structure. Indicated p130 truncation mutants were screened for their ability to be degraded following co-overexpression with cyclin F. The data supporting these conclusions are shown in Supplemental Figure S3. ***(B)*** The first and last amino acids in the two potential cyclin F binding sites in the p130 spacer domain (R658-I660 and R680-L682) were mutated to alanine (AxA). HEK293T cells transiently expressing HA-p130 alone (WT or AxA mutants, as indicated), or together with FLAG-cyclin F WT. Cells were collected and analyzed by immunoblot 24h post-transfection. (representative of n=3 experiments) ***(C)*** GST-p130^593-790^(WT) and GST-p130^593-790^(AA) were produced in *E. coli* and purified. FLAG-cyclin F was transiently expressed in HEK293T cells. GST pulldowns (left) and FLAG pulldowns (right) were used to determine the interaction of p130 and cyclin F. Binding was assessed by immunoblot after pulldown. (representative of n=3 experiments) ***(D)*** GST-p130(WT) and FLAG-cyclin F(WT) or FLAG-cyclin F(M309A L313A) were expressed as in described in *(C).* Interaction and binding were assessed as in *(C).* (representative of n=3 experiments) ***(E)*** Fluorescence polarization anisotropy assay to detect direct association of E2F1 and p130 with cyclin F. (Left) TAMRA-E2F1^84-99^ was titrated with increasing concentrations of purified GST-cyclin F^25-546^-Skp1. (Right) The E2F1 probe bound with 0.5 μM GST-cyclin F^25-546^-Skp1 was displaced with increasing concentrations of the indicated p130 protein construct or synthetic peptide (residues 674-692). Experiments were performed in triplicate, and the standard deviation is reported as the error.

Since p130(R680A L682A; hereafter referred to as p130(AA)) cannot be degraded by cyclin F, we asked whether it still binds to cyclin F. GST-tagged p130 truncations were purified from *E. coli* (amino acids 593-790; hereafter referred to as GST-p130) for both p130(WT) and p130(AA), and mixed with total lysates from HEK293T cells expressing FLAG-cyclin F. In a GST-pulldown assay, FLAG-cyclin F bound to GST-p130(WT), but not to GST-p130(AA) or GST alone (Fig. 4C, left panel). Conversely, we immunoprecipitated FLAG-cyclin F from the whole cell extract using a FLAG antibody, washed away unbound proteins, and combined this with purified GST-p130(WT) or GST-p130(AA). Similarly, cyclin F interacted with GST-p130(WT) but not GST-p130(AA) (Fig. 4C, right panel). Finally, we assessed whether the cyclin F hydrophobic patch mutant that cannot bind substrates (M309A L313A) would be able to bind GST-p130(WT). Consistent with its inability to trigger p130 degradation, cyclin F(M309A L313A) could not bind to GST-p130 (Fig. 4D). Together, these data reveal that the p130 Cy motif at R680-L682 and the cyclin F substrate-binding hydrophobic patch (M309-L313) are required for the p130-cyclin F interaction.

Next, to confirm direct binding and quantify affinity of the p130-cyclin F interaction, we developed a fluorescence polarization anisotropy assay using the Cy motif from E2F1 as a probe (Fig. 4E). We first titrated recombinant, purified cyclin F^25-546^-Skp1 dimer into solution of a synthetic E2F1 peptide (residues 84-99) containing a TAMRA label and found Kd = 0.40 ± 0.02 μM . We then competed off the labeled peptide using purified GST-p130 and GST-p130(AA) and a synthetic p130 peptide (residues 674-692) only containing the minimal Cy motif. We found that the short peptide (Ki = 0.10 ± 0.03 μM) and GST-p130(WT) (Ki = 0.44 ± 0.04 μM) bound with a similar inhibition constant, while the mutant p130 was a ∼100-fold less potent inhibitor (Ki = 28 ± 5 μM). We conclude that p130 binds cyclin F-Skp1 directly and that critical interactions are made between R680 and L682 in p130 and cyclin F. The fact that some affinity is maintained by the mutant p130(AA) suggests that some possible additional interactions involving residues beyond the RxL motif are present.

To determine p130 ubiquitination by cyclin F, we reconstituted the SCF^cyclin F^ ubiquitination reaction *in vitro* using purified components. We used a version of cyclin F, produced in baculovirus infected Sf9 cells, which lacks the first ∼25 amino acids and its C-terminal PEST domain to improve solubility (residues 25-546, cyclin F^25-546^). Cyclin F was combined with neddylated Cul1, Skp1, substrate, and ubiquitin. The human homologue of ariadne (ARIH1) is a RING-Between-RING (RBR) E3 ligase that has been shown to work with the E2 UBCH7 and other cullin ring E3 ligases, including the SCF, to ubiquitinate substrates (Horn-Ghetko et al., 2021; Scott et al., 2016). Therefore, ARIH1/UBCH7 and the chain-elongating E2 CDC34b were used as the ubiquitin transfer module.

Using this recombinant system to monitor ubiquitination, SCF^Cyclin F^ could robustly ubiquitinate GST-p130(WT) (Fig. 5A). This modification was dependent on the inclusion of Nedd8-Cul1-Roc1, Skp1-Cyclin F, or ARIH1/UBCH7 in the reaction since the exclusion of any of these components completely abrogated ubiquitination (Fig. 5A). Significantly, GST-p130(AA) was unable to be ubiquitinated by SCF^cyclin F^.

**Figure 5.**
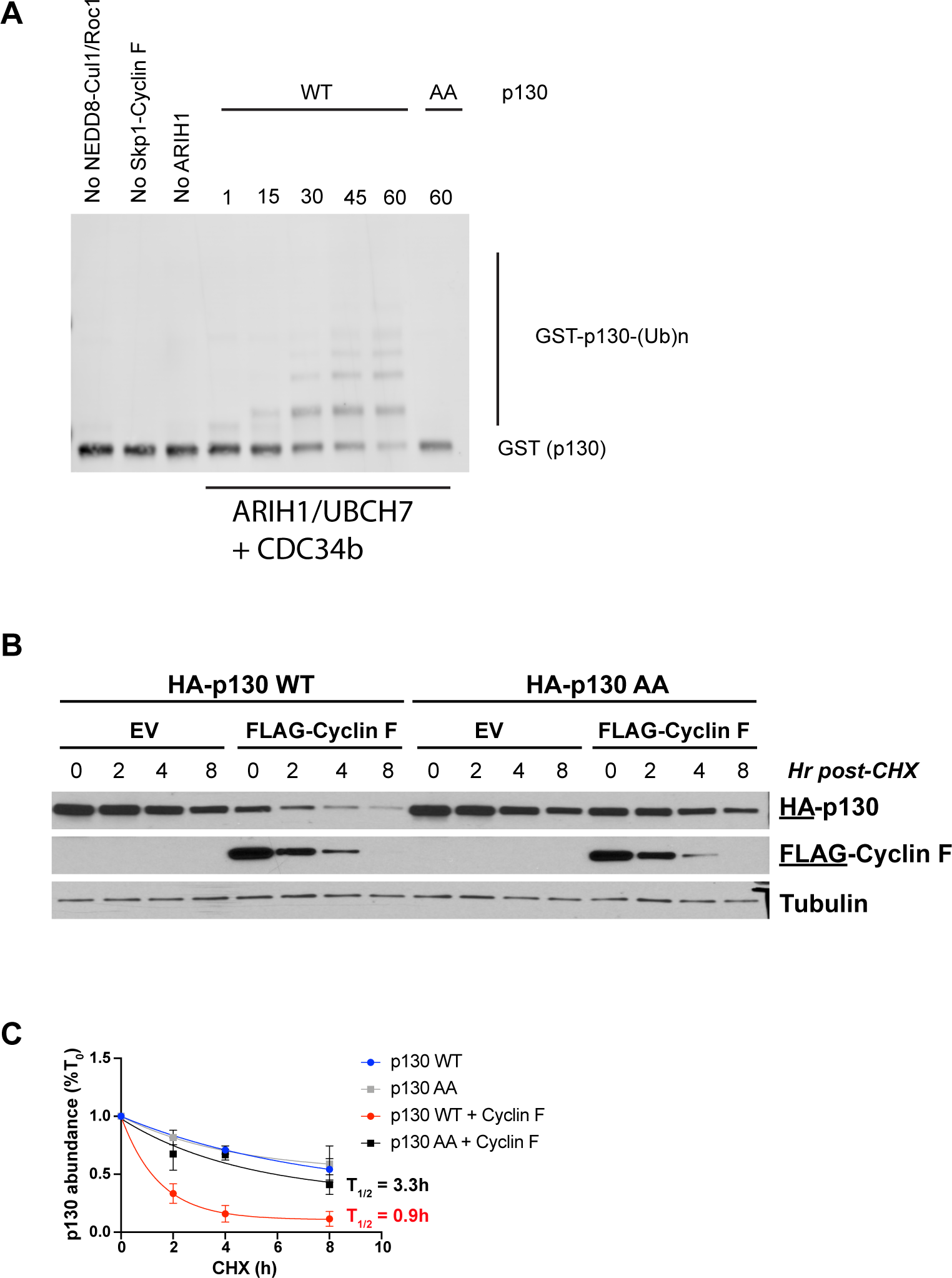
SCF^Cyclin F^ regulates the ubiquitination and stability of p130. (A) Ubiquitination reactions were performed with SCF^cyclin F^, ARIH1/Ubch7 and Cdc34b. GST-tagged p130^593-790^ WT and GST-p130^593-790^ were used as substrates and detected by immunoblot against GST. (Data are representative of n=3 experiments). (B) FLAG-cyclin F, HA-p130(WT), and/or HA-p130(AA) were transiently expressed in HEK293T cells for 24 hours. The protein synthesis inhibitor cycloheximide (CHX) was added and cells were collected at the indicated timepoints. Protein levels were determined by immunoblot. (representative of n=3 experiments) (C) Quantification of *(B).* Data is shown as mean ± SEM for n=3 experiments.

Finally, because cyclin F can ubiquitinate p130(WT), we determined the stability of p130(WT) and p130(AA) in the absence or presence of cyclin F. To determine p130 half-life, we expressed full-length HA-p130(WT) or HA-p130(AA) with or without FLAG-cyclin F for 24 hours. Cells were treated with the protein synthesis inhibitor cycloheximide (CHX) and then analyzed for HA-p130 half-life. While p130(WT) half-life was significantly reduced by ectopic expression of cyclin F, p130(AA) remained stable for the length of the experiment in the presence or absence of cyclin F (Fig. 5B-C), indicating that protein levels of p130(AA) are resistant to cyclin F-mediated degradation.

### Loss of p130 regulation by cyclin F causes G0/G1 arrest and apoptosis

Interfering with RB inactivation by inhibiting its phosphorylation, either through CDK4/6 inhibition or by expression of a mutant harboring amino acids substitution at CDK phosphorylation sites, impairs cell cycle progression (Fry et al., 2004; Lukas et al., 1997). We postulated that interfering with p130 inactivation by preventing its ubiquitination by SCF^cyclin F^ could similarly impair cell cycle and proliferation. To isolate the impact of cyclin F on p130, independent of other cyclin F substrates, we engineered NHF-1 cells with a doxycycline inducible transgene for either p130(WT) or p130(AA) (Fig. S4A). To determine whether p130(AA) accumulates more than p130(WT), we collected cells at the end of two weeks in doxycycline and immunoblotted. Although protein levels of p130(WT) and p130(AA) are initially equal (Fig. S4A), p130(AA) accumulated to higher levels compared to p130(WT) over time (Fig. 6A), consistent with p130(AA) being resistant to degradation.

**Figure 6.**
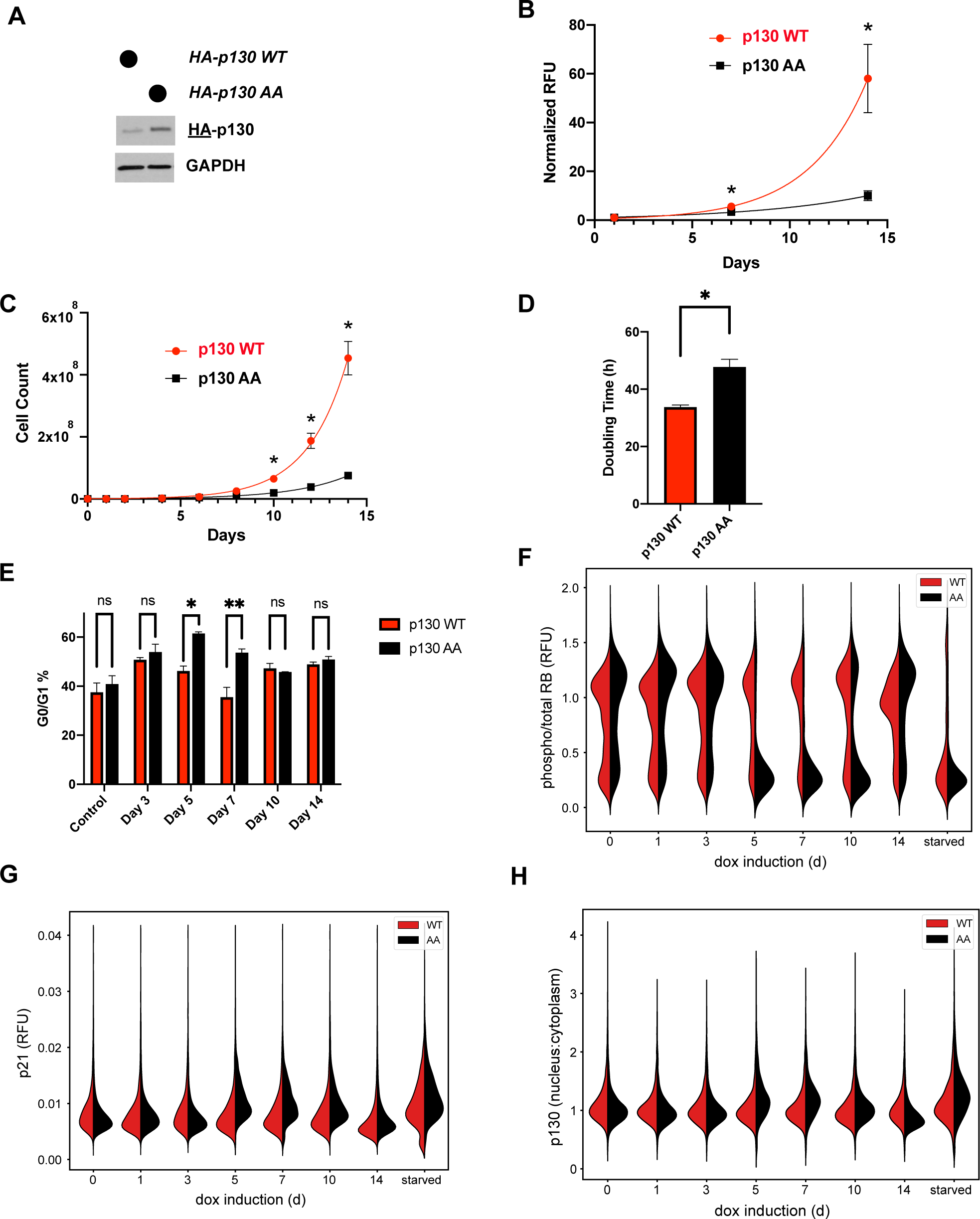
Cells expressing p130(AA) exhibit proliferation defects. ***(A)*** NHF-1 cells were engineered to express a TET-inducible HA-p130(WT) or HA-p130(AA) transgene. Doxycycline was used to induce p130 expression, and water was used as a vehicle control. HA-p130 levels were assessed by immunoblot after cells were grown in doxycycline for 14 days. ***(B)*** Inducible p130 NHF-1 cells were grown in doxycycline-containing media for 14 days to induce p130 expression. Cell number/viability was assessed on the indicated days using the PrestoBlue assay. Data represent mean ± SEM for n=3 independent experiments. ***(C)*** Inducible p130 NHF-1 cells were grown in doxycycline-containing media for 14 days to induce p130 expression. Cells were counted on the indicated days. Data represent mean ± SEM for n=3 independent experiments. ***(D)*** Doubling time was calculated from counting experiment in *(C)*. Error bars are SEM for n=3 independent experiments. ***(E)*** Inducible p130 NHF-1 cells were grown in media containing doxycycline for 14 days to induce p130 expression, or in media containing vehicle control. On indicated days, cells were pulsed with EdU for 30 minutes prior to harvest, then analyzed by flow cytometry. Data represent mean ± SEM for n=3 replicates. ***(F-H)*** Inducible p130 NHF-1 cells were grown in doxycycline for indicated times. Cells were analyzed by immunofluorescent staining and fixed cell imaging for phosphorylated vs. total RB (*F*), p21 (*G*), and nuclear vs. cytoplasmic p130 (*H*), as indicated.

To assess proliferation, we added doxycycline to induce expression of low levels of p130(WT) or p130(AA). We first used a PrestoBlue assay to indirectly assess cell number and viability based on resazurin metabolism by cells. The expression of non-degradable p130(AA) led to a profound decrease in proliferation/viability (Fig. 6B). After 14 days, the PrestoBlue fluorescence signal was decreased 5.8-fold (∼83%) in p130(AA)-expressing cells relative to controls expressing p130(WT).

Because the PrestoBlue assay suggested that cells may be growing more slowly or dying, we sought to directly quantify the proliferation of p130(WT)- and p130(AA)-expressing cells. After p130 induction in NHF-1 cells, we counted cells every 48 hours for two weeks. We found that the p130(AA)-expressing cells grew much more slowly than the p130(WT)-expressing cells (Fig. 6C). After 14 days, there were 7.7-fold (∼87%) less cells in the p130(AA) population compared to p130(WT)-expressing cells. The average doubling time for p130(WT)-expressing cells was 33h, whereas the doubling time for p130(AA)-expressing cells was increased by 45%, to 48h (Fig. 6D). In both p130(WT) and p130(AA) cell lines, the addition of vehicle alone had no observable effect on proliferation and both cell lines had an average doubling time of 27 hours across a two-week experiment (Fig. S4B-C). Expectedly, overexpression of p130(WT) alone slowed the doubling time compared to cells treated with a vehicle control, consistent with previous studies which report p130 overexpression slows cell growth (Lacy and Whyte, 1997). Nevertheless, p130(AA) had a much more severe impact on proliferation/survival. Together, these data suggest either a strong growth defect or cell death in cells expressing non-degradable p130.

We next investigated whether the slow growth observed following p130(AA)-induction resulted from cell cycle arrest. We performed flow cytometry to analyze cell cycle phase distribution using EdU incorporation and DNA staining. Inducible p130 NHF-1 cells were grown in doxycycline for 0-14 days and pulsed with EdU for 30 minutes prior to fixation. The percent of p130(AA)-expressing cells in G0/G1 significantly increased at days 5 and 7 compared to p130(WT)-expressing cells. Surprisingly, after day 10, there was no difference in G0/G1% between p130(AA) and p130(WT)-expressing cells, suggesting that the cell population may adapt to the increased levels of p130, which remain elevated throughout the entire two-week experiment (Fig. 6E).

To further examine a potential cell cycle arrest, we used iterative indirect immunofluorescence imaging (4i) to determine the levels of cell cycle markers in p130(WT)- and p130(AA)-expressing cells at various timepoints. This technique allows multiple rounds of immunofluorescence to be performed on the same population of cells(Gut et al., 2018). To determine whether cells were arresting in G1, we stained for total and phospho-RB. RB is phosphorylated in proliferating cells, while unphosphorylated RB is a hallmark of G1/G0. We found that RB phosphorylation was decreased in p130(AA)-expressing cells at days 5, 7, and 10 compared to p130(WT)-expressing cells or vehicle-treated control cells (Fig. 6F). We also found that p21 was increased in p130(AA)-expressing cells at days 5, 7, and 10, consistent with a G0/G1 arrest (Fig. 6G). To assess cell cycle distribution in each of those cell populations, we quantified total DNA content. DNA content analysis revealed that a larger percentage of p130(AA)-expressing cells had <4C DNA content compared to p130(WT)-expressing cells at days 5, 7, and 10, further indicating that the p130(AA)-expressing cells accumulate in G0/G1 phase (Fig. S5A). Finally, we assessed the subcellular localization of p130 at each timepoint by quantifying the ratio of nuclear:cytoplasmic intensity from immunofluorescence images. We found p130(AA) was more localized to the nucleus at days 5 and 7 compared to p130(WT) (Fig. 6H, S5B), consistent with p130(AA) mediating cell cycle arrest through cell cycle gene repression.

As a component of the DREAM complex, p130 functions to prevent the transcription program of myriad early (S-phase) and late (G2/M) cell cycle genes (Engeland, 2018). Because we observe an increase in G0/G1% for populations of cells expressing p130(AA) and because p130(AA) is more localized to the nucleus at those timepoints, we sought to determine the expression levels of several well-characterized DREAM target genes including: *CCNF, CCNE1, CDC6, DHFR,* and *E2F1*. To determine the RNA expression levels of these genes, we collected inducible p130(WT) or p130(AA) NHF-1 cells after 0 or 8 days in doxycycline and analyzed gene expression by RT-qPCR. There was a significant reduction in expression of *CCNF, CCNE1, CDC6, DHFR,* and *E2F1* in p130(AA)-expressing cells compared to p130(WT)-expressing cells (Fig. 7A). Since both p130(WT)-expressing cells and p130(AA)-expressing cells grew more slowly than cells not expressing a p130 transgene, we also performed RT-qPCR on control cells; the expression of *CCNF, CCNE1, CDC6, DHFR,* and *E2F1* were lower in both p130(WT)- and p130 (AA)-expressing cells compared to controls (Fig. S5E). Thus, ectopic expression of p130 reduces transcription of DREAM target genes, which is further decreased by the expression of p130(AA). To confirm that the reduction in gene expression of DREAM targets led to decreased protein expression, we again used 4i to quantify two DREAM targets by immunofluorescence: CDC6 and Cdt1. Consistent with DREAM activation, as well as observed G0/G1 arrest, protein levels of both CDC6 and Cdt1 were decreased from days 5-10 in p130(AA)-expressing cells compared to p130(WT)-expressing cells (Fig. 7B-C).

**Figure 7.**
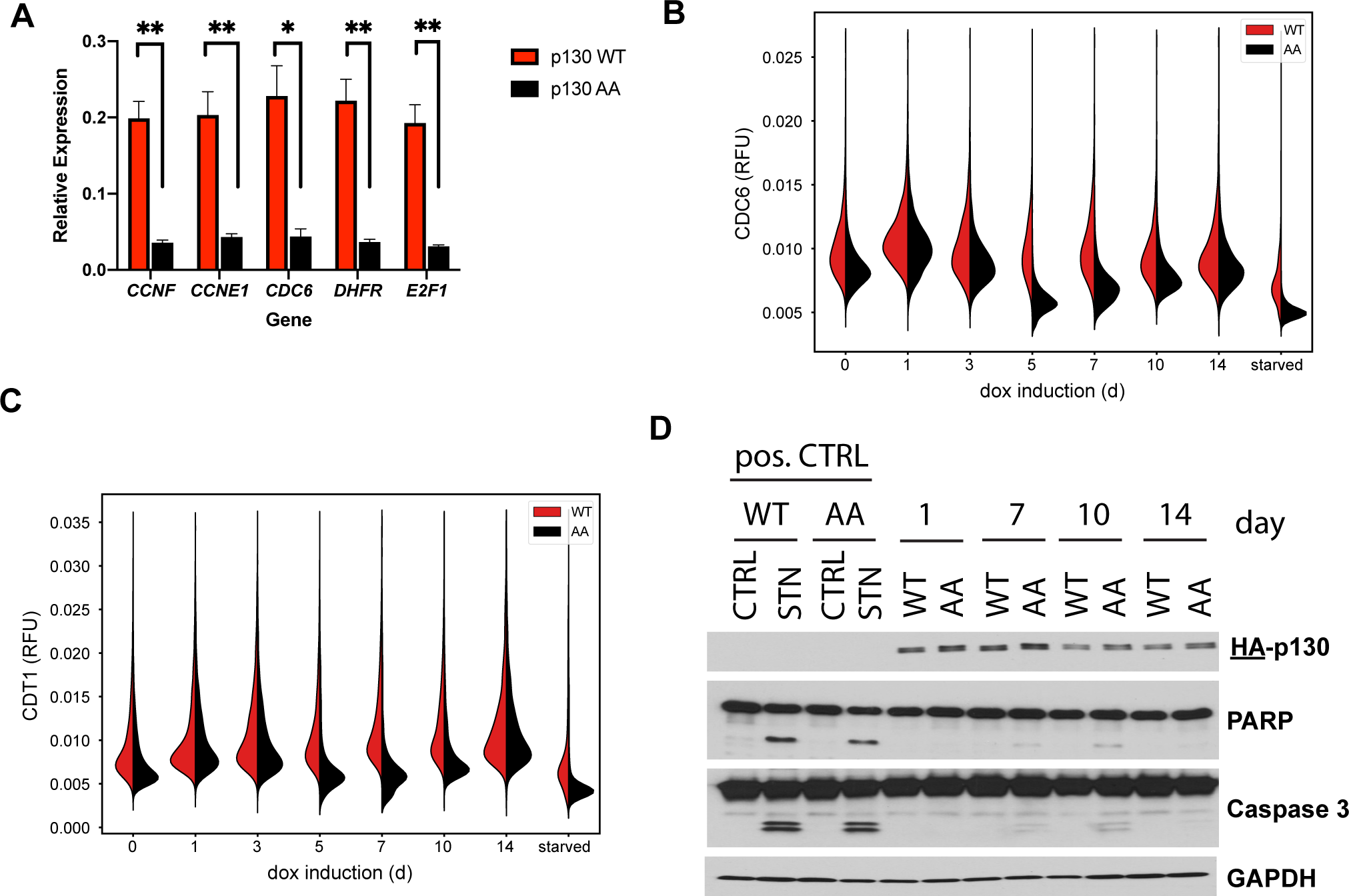
DREAM targets are downregulated, and apoptosis is induced in cells expressing p130(AA). ***(A)*** Inducible p130(WT) and p130(AA) NHF-1 cells were grown for 8 days in media containing 100 nM doxycycline to induce p130 expression. RNA was extracted for rt-qPCR analysis. Gene expression is relative to GAPDH expression and normalized to the vehicle control. Data is mean of n=3 experiments, and error bars are SEM. ***(B-C)*** Inducible p130 NHF-1 cells were grown in doxycycline for indicated times, and protein levels of CDC6 (*B*) and Cdt1(*C*) were analyzed by immunofluorescent staining and fixed cell imaging. ***(D)*** Inducible p130(WT) and p130(AA) NHF-1 cells were grown in doxycycline-containing media for the indicated number of days. Cells were collected on the indicated days and analyzed by immunoblot for the indicated proteins. As a positive control for apoptosis, cells were treated with the broad-spectrum kinase inhibitor staurosporine (or DMSO as a control). (representative of n=2 independent experiments)

Because not all cells arrest in G0/G1, despite the profound difference in cell proliferation between p130(WT) and p130(AA)-expressing cells (Fig. 6E), we asked whether another factor may be contributing to the reduced proliferation of the p130(AA)-expressing population. High p130 levels have previously been linked to apoptosis (Pentimalli et al., 2018; Ventura et al., 2018), whereas low p130 levels have been shown to protect against apoptosis (Bellan et al., 2002). To determine whether p130(AA)-expressing cells might be dying by apoptosis, we grew inducible p130(WT) and p130(AA) cells for 0-14 days in doxycycline, then immunoblotted for the apoptotic markers cleaved PARP and cleaved caspase 3. As a positive control, both cell lines were treated with DMSO or 100 nM staurosporine to induce apoptosis. We observe cleaved PARP in the p130(AA)-expressing population at days 7, 10, and 14, and cleaved caspase 3 at days 7 and 10, indicating that cells in these populations are undergoing apoptosis. We did not observe cleaved PARP or cleaved caspase in any of the populations for p130(WT)-expressing cells (Fig. 7D). Since DNA damage can cause apoptosis, and because high p21 levels can be an indicator of DNA damage, we used 4i to determine expression of proteins in the DNA damage response pathway including 53BP1, phospho-Chk1, and phospho-H2A.X. We found that in the p130(AA)-expressing cells at days 5, 7, and 10, that levels of 53BP1, phospho-Chk1, and phospho-H2A.X were downregulated (Fig. S5C-E). A decrease in DNA damage markers is consistent with fewer cells being in S-phase and suggests that DNA damage is not the cause of apoptosis in those populations. We also examined senescence but saw no apparent increase in beta-galactosidase staining in p130(AA)-expressing cells. Collectively, these results indicate that the observed growth defect caused by an inability to degrade p130 by cyclin F is a consequence of both cell cycle arrest and apoptosis.

Together, these results suggest that cyclin F contributes to the inactivation of the RB-family protein p130 and is therefore a regulator of the CDK-RB network involved in cell cycle gene repression. This is consistent with our observation that CCNF KO is highly correlated with *CDK4* and *CCND1*/*Cyclin D1* KO in the DepMap analysis. In an effort towards drug repurposing, the DepMap Consortium has tested hundreds of cell lines for their sensitivity to over 1000 oncology and non-oncology drugs. Combining these data with CRISPR/Cas9 screening data has previously been used to analyze gene-drug similarities (Corsello et al., 2020), similar to the pairwise gene-gene analyses, described above. We used these data to query the highly selective CDK4/6 inhibitors Palbociclib and Ribociclib, which are used to treat metastatic, hormone receptor positive breast cancer (Agostinetto et al., 2021; Kay et al., 2021). We determined which gene knockouts are most highly correlated with sensitivity to these CDK4/6 inhibitors. When Palbociclib was compared to data from CRISPR knockouts screens, the two most highly correlated genes are *CCND1/Cyclin D1 and CDK4,* validating this approach.

Notably, *CCNF* ranked 9^th^ among over 17,000 genes queried (Fig. S6A). For Ribociclib, *CCND1* is the most highly correlated, *CDK4* is ranked fifth, *CDK6* 52^nd^, and *CCNF* was 125^th^, falling within the top 1% of most highly correlated genes (Fig. S6B).

Conversely, we asked which drugs are highly correlated with *CDK4, CCND1, and CCNF* KO. Palbociclib and Ribociclib are among the top 4 most highly correlated drugs with *CDK4* and *CCND1* (Fig. S6C-D). Remarkably, when *CCNF/Cyclin F* knockout is compared to all drugs tested, Palbociclib is ranked first, and Ribociclib is ranked 24^th^ (Fig. S6E). Thus, across hundreds of cells lines, *CCNF/Cyclin F* KO is highly correlated with both genetic and pharmacologic inactivation of the CDK-RB network.

## Discussion

Here we demonstrate that the RB-like protein p130 is a substrate of the E3 ubiquitin ligase SCF^Cyclin F^. Cyclin F knockout or depletion allows p130 to accumulate, whereas its overexpression promotes p130 degradation. Cyclin F and p130 bind directly, and SCF^cyclin F^ can ubiquitinate p130 in a fully reconstituted *in vitro* system. We mapped a critical cyclin F binding sequence in p130, and when that sequence is mutated, it prevents p130 ubiquitination and degradation. This mutant version of p130 accumulates in normal human fibroblast cells, causing slowed proliferation, an accumulation in G0/G1 phase, and apoptosis. Together, these results tie cyclin F to the CDK-RB network and proliferative control.

### RB-family protein dynamics throughout the cell cycle

The activity of RB and the RB-like proteins, p107 and p130, oscillates during the cell cycle and quiescence. Cyclin-CDK complexes phosphorylate RB, p107, and p130, which prevents their binding to E2Fs, allowing for the transcription of E2F cell cycle target genes. In addition, RB-like proteins are also controlled at the level of their abundance. As cells exit quiescence, p130 protein is degraded by the ubiquitin-proteasome system. Then, when cells transition from proliferation to quiescence, p130 protein levels significantly increase (Smith et al., 1996). The role of phosphorylation in controlling RB and the RB-like proteins has been well-established (Canhoto et al., 2000; Farkas et al., 2002; Hansen et al., 2001), however much less is known about how changes in the levels of RB-like proteins might contribute to changes in pathway activity or its ability to restrain E2F transcription factors. Notably, as cells grow bigger, RB and its yeast ortholog Whi5, are diluted, contributing to cell cycle progression (Schmoller et al., 2015; Zatulovskiy et al., 2020). Our studies suggest that the targeted degradation of p130 by cyclin F, which effectively and rapidly reduces its concentration, also plays a significant role in promoting cell cycle progression.

Despite a concrete understanding of the role of phosphorylation in regulating RB-like proteins, less is known about the role of ubiquitin ligases. The phosphatase subunit NRBE3 was previously linked to RB ubiquitination (Wang et al., 2015). A nucleolar protein, U3 protein 14a (hUTP14a), was also suggested to be a novel type of E3 ubiquitin ligase, capable of promoting both RB and p53 degradation and cancer cell proliferation (Liu et al., 2018). However, neither NRBE3 nor hUTP14a are ubiquitin ligases. Finally, RB degradation by an unknown, Cul2-based E3 was shown in cells expressing HPV E7 oncoprotein (White et al., 2012). Previous studies have reported that SCF^Skp2^ recognizes p130 and promotes its ubiquitination (Bhattacharya et al., 2003; Tedesco et al., 2002). However, p130 still cycles in Skp2 knockout cells, suggesting additional E3 ligase(s) may be involved in p130 regulation. Surprisingly, for unknown reasons, Skp2 overexpression had no effect on p130 levels in our experiments.

### Cyclin F as a regulator of the cell cycle gene transcription

Cyclin F has previously been linked to cell cycle gene transcription via its regulation of several proteins including B-MYB, SLBP, E2F1, E2F2, E2F3a, E2F7, and E2F8 (Emanuele et al., 2020). In response to DNA damage, cyclin F is able to bind to B-MYB, and rather than ubiquitinate it, inhibits the ability of B-MYB to promote expression of mitotic genes(Klein et al., 2015). In G2, cyclin F ubiquitinates the stem-loop binding protein (SLBP), which is required during S-phase to mediate histone biogenesis (Dankert et al., 2016). Also, during G2, cyclin F catalyzes the ubiquitination of the activator E2Fs: E2F1, E2F2, and E2F3a (Burdova et al., 2019; Clijsters et al., 2019). The ubiquitination of activator E2Fs allows for their degradation, and expression of mutant E2Fs that can no longer bind to cyclin F results in increased expression of E2F target genes. Further, during G2 phase, cyclin F was shown to regulate the atypical repressor E2Fs, E2F7 and E2F8, promoting their degradation(Wasserman et al., 2020; Yuan et al., 2019). Each of these substrates implicate cyclin F in indirectly controlling cell cycle gene expression, and our study uncovers a new mechanism by which cyclin F regulates transcriptional control at the start of the cell cycle, by ubiquitinating p130.

Adding to the complexity of the role of cyclin F in cell cycle transcription, not only does cyclin F regulate p130 degradation, but p130 regulates the expression of cyclin F. Thus, cyclin F and p130 appear to exist in a double negative feedback loop, where each can downregulate the other. Interestingly, in response to DNA damage, cyclin F is down-regulated by degradation (D’Angiolella et al., 2012), and p130 has been suggested to become activated to repress cell cycle gene expression. Thus, the degradation of cyclin F in response to genotoxic stress could preserve cell cycle arrest by allowing for the accumulation of p130, while also contributing to increased nucleotide pools through the accumulation of another cyclin F substrate: RRM2 (D’angiolella et al., 2012). Determining mechanisms of cyclin F degradation in response to DNA damage, and further elucidating the potential feedback between cyclin F and p130 warrants further analysis.

### Effect of p130 overexpression on proliferation

We have shown that expressing the mutant p130(AA), but not p130(WT), causes growth arrest and apoptosis. These results are consistent with clinical observations of outcomes of p130 expression levels in cancerous tissues. We observed that when p130 is non-degradable by cyclin F, it accumulates in cells, causing a decrease in cellular proliferation. In lung cancers, p130 expression negatively correlates with histological grading and metastasis (Alfonso Baldi, Vincenzo Esposito, Antonio De Luca, Yan Fu, Ilernando Meoli et al., 1997). For endometrial cancer patients who receive surgery, low p130 levels are significantly associated with increased recurrence and death (Susini et al., 1998). In vulvar carcinomas, loss of p130 and p27 has been shown to contribute to carcinogenesis (Zamparelli et al., 2001). In oral squamous cell carcinoma, p130-negative cases have worse prognoses than p130-positive cases (Tanaka et al., 2001). And, finally, in thyroid neoplasms, reduced p130 expression has been linked to the aggressive characteristics of anaplastic carcinoma, while high p130 expression in micropapillary carcinoma has been linked to the smaller size of those tumors (Ito et al., 2003). Thus, cancers with low p130 expression or p130 loss, seem to grow more quickly and aggressively, whereas cancers with high p130 expression are correlated with better prognoses. Our findings fit within the context of these studies; as p130(AA) accumulates, the cells are less proliferative and some die by apoptosis. Since p130 mRNA levels are not regulated during the cell cycle, it is interesting to speculate that enhancing p130 degradation in some malignancies could contribute to defects in regulation of cell cycle gene expression and hyper-proliferation. Likewise, blocking p130 degradation could phenocopy CDK4/6 inactivation and could potentially be therapeutically advantageous, and would be consistent with our analysis showing that loss-of-function in cyclin D, CDK4 and cyclin F are highly correlated across nearly 800 cancer cell lines.

Despite the growth arrest and apoptotic phenotype, the p130(AA)-expressing cell population seems to recover by two weeks of p130 overexpression. While differences were observed at earlier timepoints, at day 10 and beyond, the p130(AA) and p130(WT)-expressing populations are not significantly different in the proportion of cells in G0/G1. Further, we noticed that levels of apoptotic markers in the p130(AA)-expressing population are lower at day 14 than at earlier days. It is possible that only a subpopulation of cells is affected by the p130(AA) expression, and then those cells die by apoptosis. Alternatively, all cells could be affected by p130(AA) expression, some die, but most recover and adapt to the increase in p130 expression. Why some cells are more disadvantaged than others remains unknown. If all p130(AA)-expressing cells are affected and they overcome the growth arrest, the question remains of how they have achieved adaptation, either through evasion of apoptosis or reduced growth arrest.

Cyclin F is a strongly cell cycle regulated gene. Cyclin F mRNA levels peak in G2/M, and no studies have identified *CCNF* as an E2F target at G1/S. It is therefore unclear how its protein levels increase early in the cell cycle, in lockstep with Skp2, a bona fide E2F target gene. We previously showed that AKT can phosphorylate and stabilize cyclin F, and that this could contribute to its accumulation near G1/S (Choudhury et al., 2017). Thus, the activation of cyclin F by AKT could potentially lead to the degradation of p130. Given the recurrent activation of the PI3K-AKT pathway in many cancers, it will be interesting in the future to determine if cyclin F phosphorylation promotes p130 degradation. This would suggest that hyper-activated PI3K, either through mutation of PIK3CA or loss of PTEN, functions through cyclin F to render p130 inactive via the ubiquitin pathway.

## Acknowledgements

We thank lab members for helpful discussions throughout this project. We thank Brenda Schulman (Max Planck Institute of Biochemistry) for generously providing SCF complex reagents used for *in vitro* ubiquitination assays. We thank Larissa Litovchick (Virginia Commonwealth University) for providing the pBABE-p130 vector. We thank Dennis Goldfarb (Washington University) for help with DepMap data downloads and analysis. The Emanuele lab (TPE and MJE) is supported by the UNC University Cancer Research Fund (UCRF), the National Institutes of Health (R01GM120309, R01GM134231), and the America Cancer Society (Research Scholar Grant; RSG-18-220-01-TBG). The Brown lab (ETW and NGB) is supported by UCRF and National Institutes of Health (R35GM128855) and ETW is partially supported by R01CA163834. The Rubin lab (PN and SMR) is supported by the National Institutes of Health (R01GM127707). The Purvis lab (WMS and JEP) is supported by by NIH grants R01-GM138834 (JEP), DP2-HD091800 (JEP), and NSF CAREER Award 1845796 (JEP).

## Methods

### Acquisition and Analysis of DEPMAP data

Gene co-dependencies were determined from the Achilles dataset from depmap.org (Achilles_gene_effect.csv, downloaded 7/19/19). The Achilles dataset contains dependency scores from genome-scale essentiality screens scores of 789 cell lines. As a measure of co-dependency, the Pearson’s correlation coefficient of essentiality sores was computed for all gene pairs. Gene ontology analysis for the top 100 genes co-dependent with CCNF was performed using MetaScape. Potential Cyclin F substrates were identified by proteins encoded by genes that have a Pearson’s correlation coefficient of >0.15 when compared to CCNF and are classified by the GO term: cell division (0051301).

### Cell Culture

All cell lines were cultured in Dulbecco’s Modified Eagle’s Medium (DMEM; GIBCO) supplemented with 10% Fetal Bovine Serum (FBS; VWR) and 1% Pen/Strep (GIBCO). Cells were incubated at 37°C, 5% CO2.

DNA transfection experiments were performed in HEK293T cells for 24 hours using either Lipofectamine 2000 (ThermoFisher) or PolyJet (SignaGen) transfection reagents according to the manufacturer’s protocol. RNA transfection experiments were performed in NHF-1 and IMR-90 cells for 48 hours using RNAiMAX (ThermoFisher) transfection reagent according to the manufacturer’s protocol. Plasmid and siRNA information is located in Table S1.

To produce lentivirus, HEK293T cells were transfected with pLV[Exp]-CMV>Tet3G/Hygro, pLV[TetOn]-Neo-TRE3G>HA/{p130 WT}, or pLV[TetOn]-Neo-TRE3G>HA/{p130 AA}, and the lentivirus packaging plasmids VSV-G, Gag-pol, Tat, and Rev. Media containing virus particles was collected after 48 hours, filtered with a 0.45 µM filter, and frozen at -80°C. NHF-1 cells were transduced with a 1:1 ratio of fresh media plus lentivirus-containing media. NHF-1 cells were first transduced with the Tet3G plasmid and selected in media containing 50 µg/mL hygromycin. Then NHF-1 cells stabl expressing Tet3G were infected with either p130 WT or p130 AA and selected in media containing 375 µg/mL G418.

### Cloning and Directed Mutagenesis

pDEST-HA3-p130, pDEST-HA3-p130^1-417,^ pDEST-HA3-p130^418-1139^, pDEST-HA3-p130^418-616^, pDEST-HA3-p130^418-827^, pDEST-HA3-p130^418-1024^, pDEST-HA3-p130^828-1139^, and pINDUCER20 CCNF were produced using Gateway Cloning Technology. pLV[Exp]-CMV>Tet3G/Hygro, pLV[TetOn]-Neo-TRE3G>HA/{p130 WT}, or pLV[TetOn]-Neo-TRE3G>HA/{p130 AA} were all ordered from VectorBuilder. pGEX-GST-p130^593-790^ was a gift from Peter Whyte (McMaster University) and pINDUCER20 was a gift from Stephen Elledge (Addgene #44012). pDEST-HA3-p130 R658A I660A, pDEST-HA3-p130 R680A L682A, and pGEX-GST-p130^593-790^ R680A L682A were generated by site-directed mutagenesis using the Q5 Site-Directed Mutagenesis Kit (NEB) according to the manufacturer’s instructions. For recombinant expression, Cyclin F^25-546^ was subcloned into a pFastBac vector with an N-terminal GST tag and TEV protease site. The Skp1 gene (a gift from Dr. Bing Hao) was subcloned into a PGEX-4T-1 vector previously engineered to contain a TEV protease site. Sequences for all primers are included in Table S1.

### Cell lysis and Immunoblotting

Cells were lysed on ice for 10 min in NETN (20 mM Tris pH8.0, 100 mM NaCl, 0.5 mM EDTA, 0.5% NP40) supplemented with 10 µg/mL aprotinin, 10 µg/mL leupeptin, 10 µg/mL pepstatin A, 1 mM sodium orthovanadate, and 1 mM AEBSF(4-[2 aminoethyl] benzenesulfonyl fluoride).

Following incubation, cells were spun at 20,000 xg in a benchtop microcentrifuge at 4°C for 10 min. Protein concentration was determined by Bradford assay (Bio-Rad) and samples were prepared by boiling in Laemmli buffer. Protein was separated by electrophoresis on either homemade or TGX (Bio-Rad) stain-free gels and then was transferred to nitrocellulose membranes. Blocking was performed in 5% non-fat dry milk (Blotting Grade blocker; Bio-Rad) diluted in TBS-T (137 mM NaCl, 2.7 mM KCl, 25 mM Tris pH 7.4, 1% Tween-20). All primary antibody incubations were carried out overnight, rocking, at 4°C, and all HRP-conjugated antibody incubations were carried out for 1h, rocking, at room temperature. TBS-T was used for all wash steps. Protein abundance was visualized by chemileminescence using Pierce ECL (ThermoFisher). A detailed list of antibodies and dilutions is included in Table S1.

### Endogenous Immunoprecipitation

Asynchronous HEK293T cells were lysed and total protein was quantified as described above. 10% of the protein sample was retained as the input, and the remaining protein was divided in half to be incubated with either rabbit IgG control or Cyclin F antibodies for IP. Cell lysates were incubated for 6 hours, rotating, at 4°C with 1µg antibody per mg protein. During the incubation, Pierce Protein A/G agarose beads (ThermoFisher) were washed 3X 20 min in NETN, then resuspended at a 50% slurry in NETN. 40 µL slurry per 1 mg protein was added to the protein/antibody IP mix and incubated for 45 min, rotating, at 4°C. Beads were then washed 3X 5 min in NETN, and finally re-suspended in 2X Laemmeli buffer and boiled. Co-IP was assessed by immunoblotting as described.

### Protein Purification

In order to purify GST and GAT-p130 for the *in vitro* binding assay, BL21 *E. coli* were transformed with GST, GST-p130^WT^ or GST-p130^AA^ and a 0.5 L culture was grown to an O.D. of 0.6 then induced with 1 mM IPTG (Isopropyl β-D-1-thiogalactopyranoside) for 4 hours. Cells were harvested by centrifugation and incubated for 20 min in NTEN *E. coli* Lysis buffer (100 mM NaCl2, 20 mM Tris pH 8.0, 1 mM DTT, 1 mM EDTA, 1% NP40) supplemented with 10 µg/mL aprotinin, 10 µg/mL leupeptin, 10 µg/mL pepstatin A, and 10 mg/mL lysozyme. Then, cells were lysed by sonication at 50% power for 3X 30 second pulses, with 1 minute on ice between pulses. Lysates were then spun at 30,000 xg for 30 min at 4°C to clarify. During the clarification step, glutathione agarose resin (GoldBio) was equilibrated 3X 10 min in NTEN *E. coli* Lysis buffer. 250 µL of equilibrated glutathione agarose resin was then added to the clarified lysate and incubated for 2 h, rotating, at 4°C. Beads were washed 3X 5 min in NTEN *E. coli* Lysis buffer. Bound GST-p130 was eluted by three sequential elution steps: 2X 30 min and one overnight incubation, rotating, at 4°C in elution buffer (50 mM Tris pH 8.0, 5 mM reduced glutathione). Elution fractions were pooled, and buffer exchange into 50 mM Tris was performed by 5X buffer exchange in 30,000 MWCO spin columns.

For *in vitro* ubiquitination assays, UBA1, CDC34b, UBCH7, ARIH1, Cul1-Rbx1, and ubiquitin were expressed and purified as described previously(Kamadurai et al., 2013; Scott et al., 2016). The neddylation of Cul1-Roc1 was also performed as described previously (Duda et al., 2008). GST-p130^wt^ (residues 593-790) and GST-p130^AA^ (residues 593-790 R680A L682A) fusions were expressed in *E. coli* BL21-CodonPlus (DE3)-RIL, and then purified by glutathione-affinity and size-exclusion chromatography.

For Cyclin F^25-546^, 1.2 L of Sf9 cells were infected with baculovirus at a density of 2 x 10^6^ cells/mL of culture and harvested after 3 days. Pelleted cells were resuspended in lysis buffer (25 mM Tris, 200 mM NaCl, 1 mM DTT, 1 mM PMSF, 1x protease inhibitor cocktail, pH 8.0) and passed through a C3 emulsiflex homogenizer (AVESTIN, Inc.). After lysate clarification, the supernatant was loaded onto a GS4B glutathione agarose (Cytvia) column. The column was washed in lysate buffer lacking protease inhibitors and eluted in the same buffer with 10 mM glutathione. Purified TEV protease (1% by mass) was added overnight while protein was dialyzed in buffer containing 25 mM Tris, 200 mM NaCl, and 1 mM DTT (pH 8.0). Protein was then passed back through a GS4B column to remove free GST, concentrated, and loaded in 1 mL onto a Superdex 200 column (Cytvia) equilibrated in similar buffer. Peak fractions from the column elution were pooled and protein concentrated to 8 μM.

Full length Skp1 was expressed in E. coli as a GST fusion protein. 6L of BL21(DE3) cells were grown to O.D. of 0.6 then induced with 1 mM IPTG (Isopropyl β-D-1-thiogalactopyranoside) for 16 hours at room temperature. Protein was purified similar to as described above for cyclin F^25-546^, except that following the initial GS4B elution, protein was further purified by Source Q Sepharose ion exchange chromatography prior to TEV cleavage. For the fluorescence polarization anisotropy assays, the GST tag was left on the cyclin F for stability, and the fusion protein was mixed with Skp1 prior to the experiment.

### p130-Cyclin F *in vitro* binding assays

GST pulldowns were performed by loading 1 µg of purified GST or GST-p130 onto 15 µL of glutathione beads (GoldBio) in NETN (20 mM Tris pH8.0, 100 mM NaCl, 0.5 mM EDTA, 0.5% NP40) for 1 h, rotating at 4°C. Loaded beads were washed 3X 5 min in NETN, then incubated with 0.5 mg of whole cell extract of HEK293T cells transiently transfected with a FLAG-Cyclin F plasmid for 2h, rotating, at 4°C. Beads were again washed 3X 5 min in NETN and p130-cyclin F interaction was assessed by immunoblot.

FLAG pulldowns were performed by immunoprecipitating FLAG-Cyclin F from whole cell extracts of HEK293T cells transiently transfected with FLAG-Cyclin F. IP was performed by incubating 1 mg whole cell extract with 50 µL EZView Red anti-FLAG M2 affinity gel (MilliporeSigma) for 2 h, rotating, at 4°C. Loaded affinity gel was washed 3X 5 min with NETN, then incubated with 1 ug purified GST or GST-p130 for 2 h, rotating, at 4°C. Loaded affinity gel was again washed 3X 5 min in NETN and p130-cyclin F interaction was assessed by immunoblot.

### *In vitro* Ubiquitination Assay

Multiple turnover reactions were set up by combining UBA1, MgATP, CDC34b, HHARI, UBCH7, Nedd8∼Cul1-Roc1, SKP1, Cyclin F^25-546^, and either GST-p130 or GST-p130^AA^ in the assay buffer (20mM HEPES pH=8, 200mM NaCl) while kept on ice. The mixtures were then equilibrated to room temperature and the reaction was started by the addition of ubiquitin.

Reactions were quenched by adding SDS loading buffer at specified time points. After SDS-PAGE, the ubiquitination of GST-p130 was monitored by western blot analysis using a-GST antibody (SC-138, mouse) and a fluorescent secondary antibody (goat anti-mouse IgG (H+L), Alexa Fluor 633, A-21052). Fluorescence was observed using the Amersham Typhoon 5.

### Proliferation Assays

Cell counting: NHF-1 inducible p130 cells were seeded in 10 cm plates, and p130 WT or AA was induced by the addition of 100 nM doxycycline (or DMSO as a control) to media. Cells were trypsinized (Gibco), counted with an automatic cell counter (Bio-Rad), and replated in fresh doxycycline-containing media every 48 hours for two weeks. Cell count was plotted vs. time, differences were assessed using a student’s t-Test, and cell count was used to calculate doubling times. Experiments were carried out in triplicate and repeated three times.

PrestoBlue: NHF-1 inducible p130 cells were seeded in 96-well plates, and p130 WT or AA was induced by the addition of 100 nM doxycycline to media. Media was replaced with fresh doxycycline-containing media every 48 hours for two weeks, and growth was assessed with the PrestoBlue cell viability reagent (Thermo Fisher) on days 1, 7, and 14. Experiments were carried out with six technical replicates, and repeated three times.

Crystal Violet Assay: NHF-1 inducible p130 cells were seeded in 60 mm plates, and p130 WT or AA was induced by the addition of 100 nM doxycycline (or DMSO as a control). Media was replaced with fresh doxycycline-containing media (or DMSO-containing media as a control) every 48 hours for 10 days. On day 10, confluence was visualized by staining with Crystal Violet Staining Buffer (0.5% crystal violet, 20% methanol) for 15 minutes at room temperature, and de-staining with water. Experiments were repeated three times.

### RT-qPCR

In NHF-1 inducible p130 cells, p130 WT or AA was induced by the addition of 100 nM doxycycline (or DMSO as a control) for 14 days. Cells were harvested and RNA was extracted using the RNeasy plus mini kit (QIAGEN). 1 µg of extracted RNA was used to generate cDNA libraries using the SuperScript III First-Strand synthesis system (Thermo Fisher) following manufacturer’s instructions. Samples were diluted 1:10, except for GAPDH analysis that were diluted 1:1000. Transcript abundance was quantified using the SSO Advanced Universal SYBR Green Supermix (Bio-Rad) and measured with a QuantStudio 6 Flex Real-Time PCR system (Thermo Fisher), and transcript levels were normalized to GAPDH. Relative quantity of transcripts was quantified using the 2-ΔΔCT method. Each sample was run in triplicate. Primers are listed in Table S1.

### Flow Cytometry for Cell Cycle Analysis

In NHF-1 inducible p130 cells, p130 WT or AA was induced by the addition of 100 nM doxycycline (or DMSO as a control) for the indicated number of days. 30 minutes prior to fixation, cells were pulsed with 10 µM EdU and were fixed in 4% formaldehyde/PBS for 15 min at room temperature. Cells were pelleted and re-suspended in 1% BSA/PBS. EdU was labeled with Alexa Fluor 488 using click chemistry as previously described (CITE JEN’s PAPER). Flow was carried out on an Attune NxT Flow Cytometer (Thermo Fisher), and data was analyzed using FlowJo software.

### Florescence Polarization Anisotropy Assay

For the GST-cyclin F^25-546^-Skp1 titration, a TAMRA-labeled E2F1^84-99^ synthetic peptide at 10 nM was mixed with varying concentrations of protein complex in a buffer containing 25 mM Tris, 150 mM NaCl, 1 mM DTT, 0.1% (v/v) Tween-20, pH 8.0. For the Ki measurements, varying concentrations of GST-p130^593-790^ WT, GST-p130^593-790^ R680A/L682A mutant, or synthetic p130^674-692^ peptide were mixed with 10 nM TAMRA-labeled E2F1^84-99^ and 0.5 μM GST-cyclin F-Skp1 in a buffer containing 25 mM Tris, 150 mM NaCl, 1 mM DTT, 0.1% (v/v) Tween-20, pH 8.0. Forty microliters of the reaction were used for the measurement in a 384-well plate.

Fluorescence anisotropy (FA) measurements were made in triplicate, using a Perkin-Elmer EnVision plate reader. The KD and Ki values were calculated using Prism 8 (Version 8.4.3).

### Iterative indirect immunofluorescence imaging (4i)

Cells were plated in glass-bottom plates (Cellvis) treated as required and prepared as follows. In between each step, samples were rinsed 3X times with phosphate-buffered saline (PBS) and incubations were at room temperature, unless otherwise stated. Cells were fixed with 4% paraformaldehyde (ThermoFisher Scientific, 28908) for 30 min, permeabilized with 0.1% Triton X-100 in PBS for 15 min and inspected for sample quality control following Hoechst staining (1:1000, Millipore Sigma, 99403) in imaging buffer (IB: 700 mM N-acetyl-cysteine (Sigma, A7250) in ddH2O. Adjust to pH 7.4). Cells were rinsed 3X with ddH2O and incubated with elution buffer (EB: 0.5M L-Glycine (Sigma, 50046), 3M Urea (Sigma, U4883), 3M Guanidine chloride (ThermoFisher Scientific, 15502-016), and 70mM TCEP-HCl (Sigma, 646547) in ddH20.

Adjusted to pH 2.5) 3X for 10 min on shaker to remove Hoechst stain. Sample was incubated with 4i blocking solution (sBS: 100 mM maleimide (Sigma, 129585), 100 mM NH4Cl (Sigma, A9434) AND 1% bovine serum albumin in PBS) for 1h and incubated with primary antibodies (Table S1) diluted as required in conventional blocking solution (cBS: 1% bovine serum albumin in PBS) overnight at 4°C. Samples were rinsed 3X with PBS and then incubated in secondary antibodies (1:500, donkey anti-rabbit AlexaFluor Plus 488 (ThermoFisher Scientific, A32790), donkey anti-mouse AlexaFluor Plus 555 (ThermoFisher Scientific, A32773) and donkey anti-goat AlexaFluor Plus 647 (ThermoFisher Scientific, A32758)) and Hoechst for 1h on shaker, then rinsed 5X with PBS and imaged in IB. Samples were imaged using the Nikon Ti Eclipse inverted microscope with a Nikon Plan Apochromat Lambda 40x objective with a numerical aperture of 0.95 and an Andor Zyla 4.2 sCMOS detector. Stitched 8x8 images were acquired for each condition using the following filter cubes (Chroma): DAPI(383-408/425/435-485nm), GFP(450-490/495/500-550nm), Cy3(530-560/570/573-648nm), Cy5(590-650/660/663-738nm).

After imaging, samples were rinsed 3X with ddH2O, antibodies were eluted and re-stained iteratively as described above. Nuclear/cytoplasmic segmentation and quantification was performed using standard modules in CellProfiler (v3.1.8) and data was visualized using a custom Python script (v3.7.1)

### Quantification and Statistical Analysis

Western blot quantification was carried out using the Fiji software. All statistical analyses were carried out using GraphPad Prism v9.

## Supplemental Figure Legends

**Figure S1.**
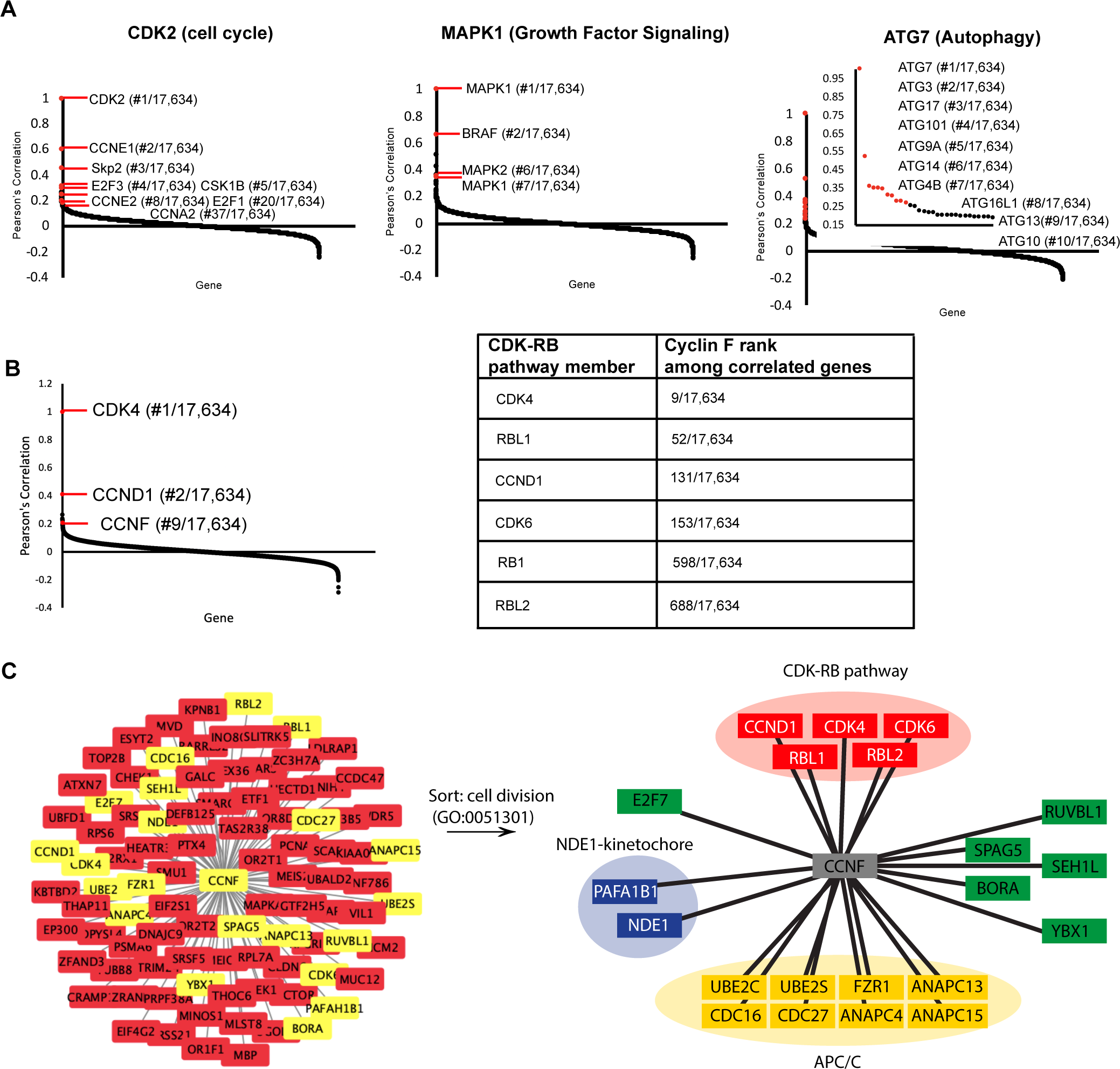
Analysis of Project Achilles Dependency Map reveals that *CDK-RB* network genes correlate highly with *CCNF*. ***(A)*** Cancer Dependency Map data from Project Achilles was analyzed to identify genes whose loss-of-function impact on cellular fitness is correlated with known proliferative regulators. Data are based on pooled CRISPR/Cas9 gene knockout screens performed in 789 cell lines. Pearson’s correlation coefficients for genes correlated with *CDK2* (left), *MAPK1* (middle) and *ATG7* (right). Known interactors are highlighted in red. Pearson’s correlation coefficients are reported for all gene pairs (each dot corresponds to a gene pair). ***(B)*** Pearson’s correlation coefficients for genes whose fitness score correlates with *CDK4*. Genes of interest are highlighted in red (left). *CCNF* rank among genes correlated with CDK4, *CDK6, CCND1, RB1, RBL1,* and *RBL2* (right). ***(C)*** The top 94 genes whose fitness score is most highly correlated with CCNF (Pearson correlation greater than 0.15) were filtered by the GO term cell division (GO:00513010). All 94 are shown on the left, with those that are classified under the GO term cell division highlighted in yellow. The 21 sorted genes fall into four categories: NDE1-kinetochore complex (blue), CDK-RB pathway (red), Anaphase promoting complex/Cyclosome (yellow), and other (green). This map is the same as that shown in Figure 1.

**Figure S2.**
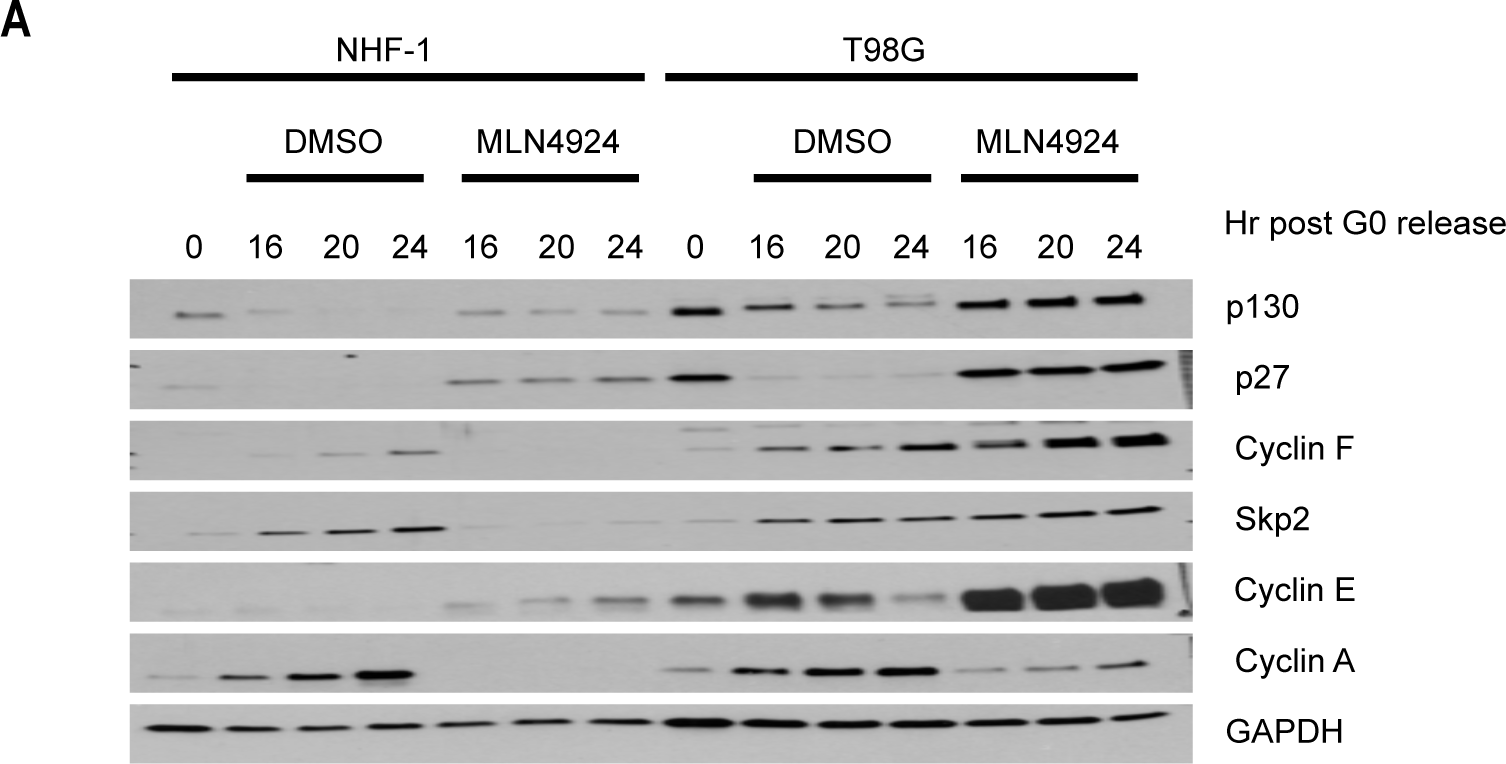
p130 dynamics in MLN4924-treated cell lines exiting quiescence. ***(A)*** NHF-1 and T98G cells were synchronized in G0 by serum starvation for 48 h. Cells were then released into the cell cycle upon the addition of serum-containing media supplemented with either MLN4924 or vehicle (DMSO) as a control. Cells were collected at the indicated time points after release. Protein levels were assessed by immunoblot. (Data represent n=2 independent experiments for T98G cells and n=3 independent experiments for NHF-1 cells)

**Figure S3.**
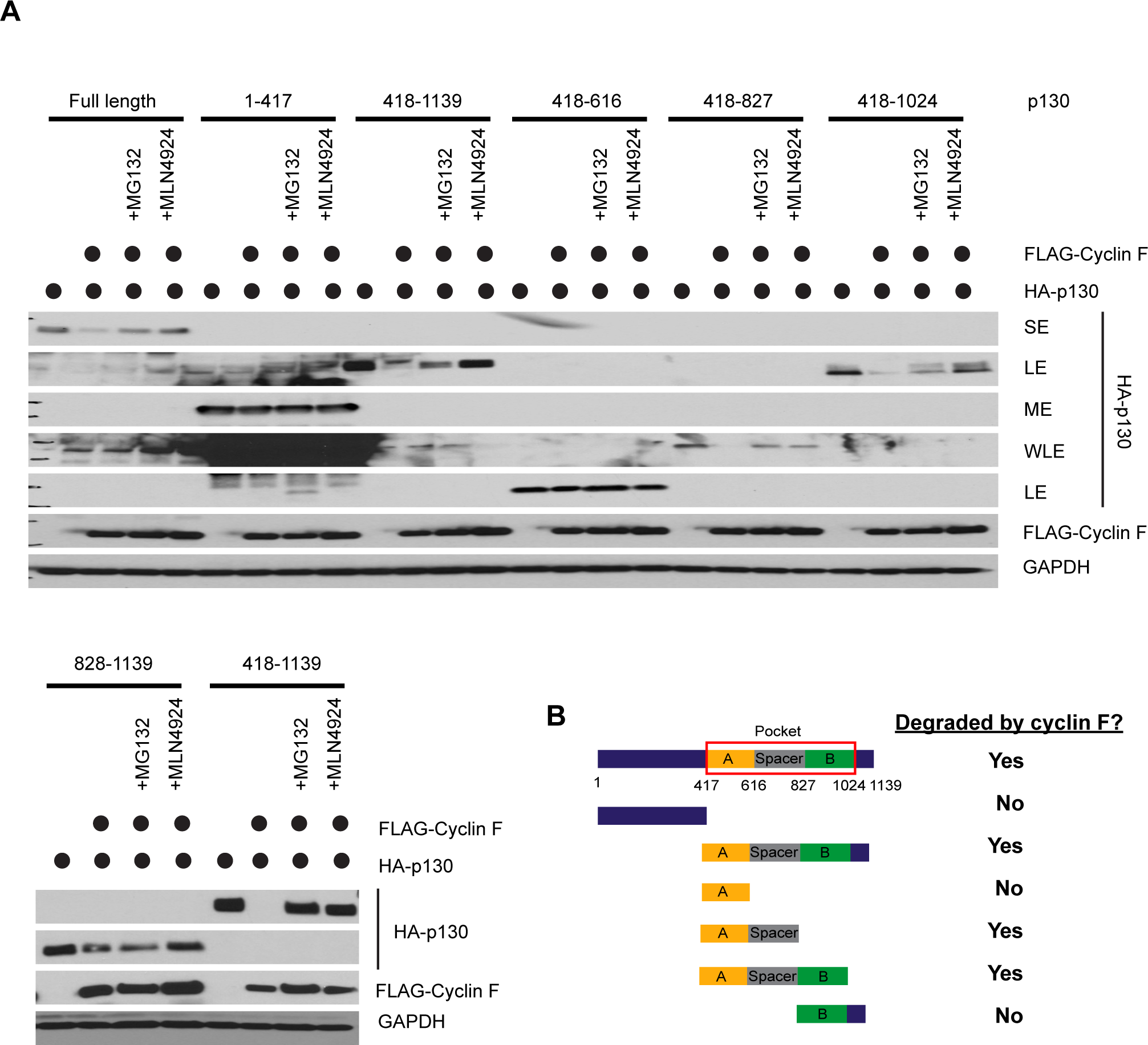
Degradation of p130 truncation mutants following cyclin F co-expression requires the p130 spacer region. ***(A)*** HEK293T cells were transiently transfected for 24 hours with the indicated HA-tagged p130 truncations either alone or in combination with FLAG-cyclin F. MG132 or MLN4924 were added for the last six hours where indicated. Cells were collected 24-hours post transfection and analyzed by immunoblot. ***(Top)*** p130 full-length (amino acids 1-1139), p130^1-417^, p130^418-1139^, p130^418-616^, p130^418-827^, and p130^418-1024^ and ***(Bottom)*** p130^828-1139^ and p130^418-1139^. (Blots are representative of n=3 replicate experiments. SE=short exposure, ME=medium exposure, LE=long exposure, WLE=wicked long exposure) ***(B)*** Graphical depiction of p130 domain structure and truncations assessed in *(A)*.

**Figure S4.**
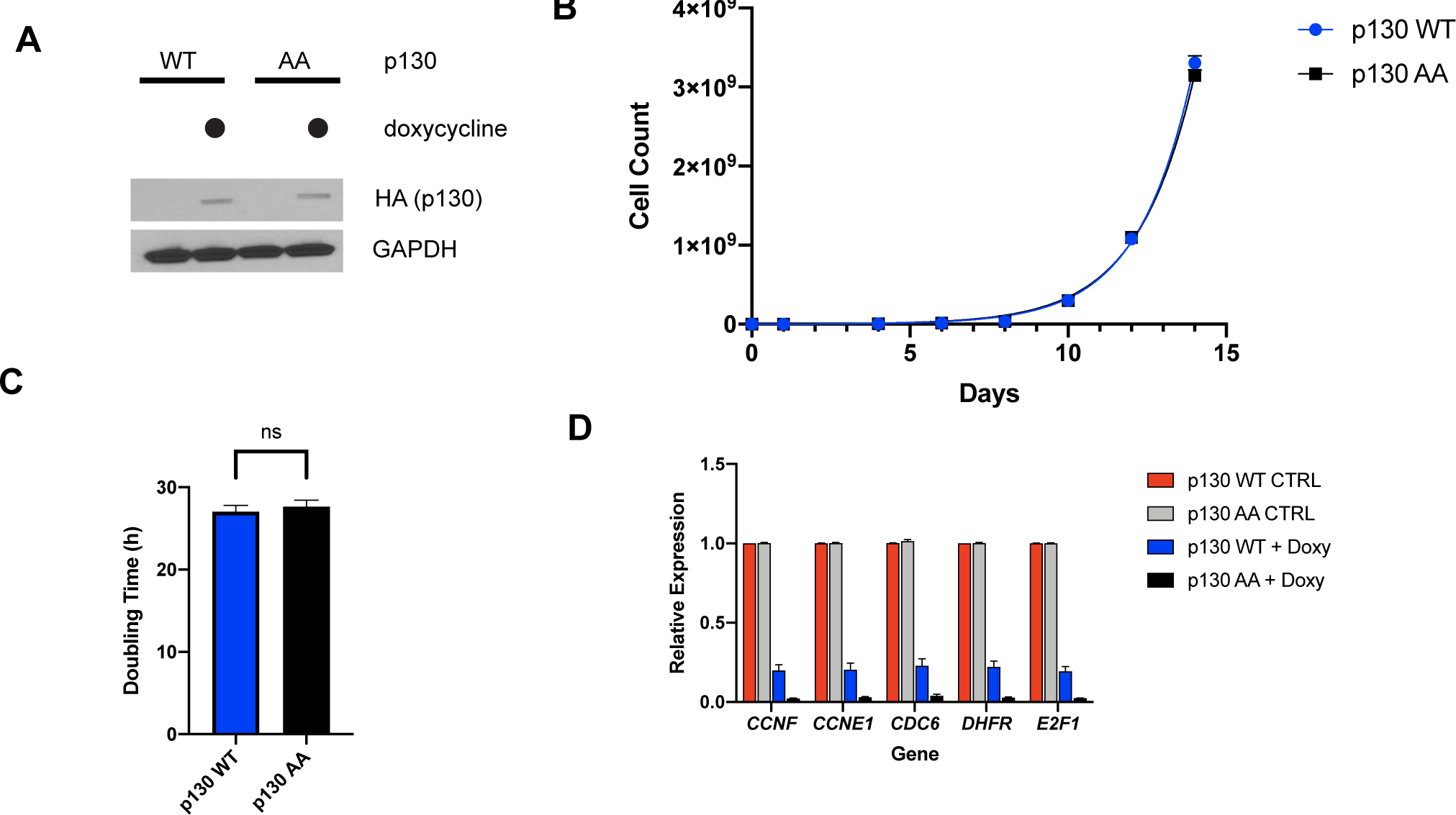
TET-inducible p130 WT and p130 AA NHF-1 cells grow proliferate rapidly prior to induction of p130 expression. ***(A)*** p130(WT) and p130(AA) expression is induced with 100 nM doxycycline for 24 hours. No p130 expression is detected in vehicle control (water) inductions. ***(B)*** Inducible p130(WT) and p130(AA) NHF-1 cells were grown for two weeks in media containing vehicle (water) as control induction of p130. Cells were counted at the indicated times. Data represent mean of n=3 experiments ± SEM. ***(C)*** Doubling time was calculated from counting data in *(B)*. Error bars are SEM. ***(D)*** Inducible p130(WT) and p130(AA) NHF-1 cells were grown in for 8 days in media containing 100 nM doxycycline to induce p130 expression (or containing water as a control). RNA was extracted for rt-qPCR analysis. Gene expression is relative to GAPDH expression and normalized to the vehicle control. Data is mean of n=3 experiments and error bars are SEM.

**Figure S5.**
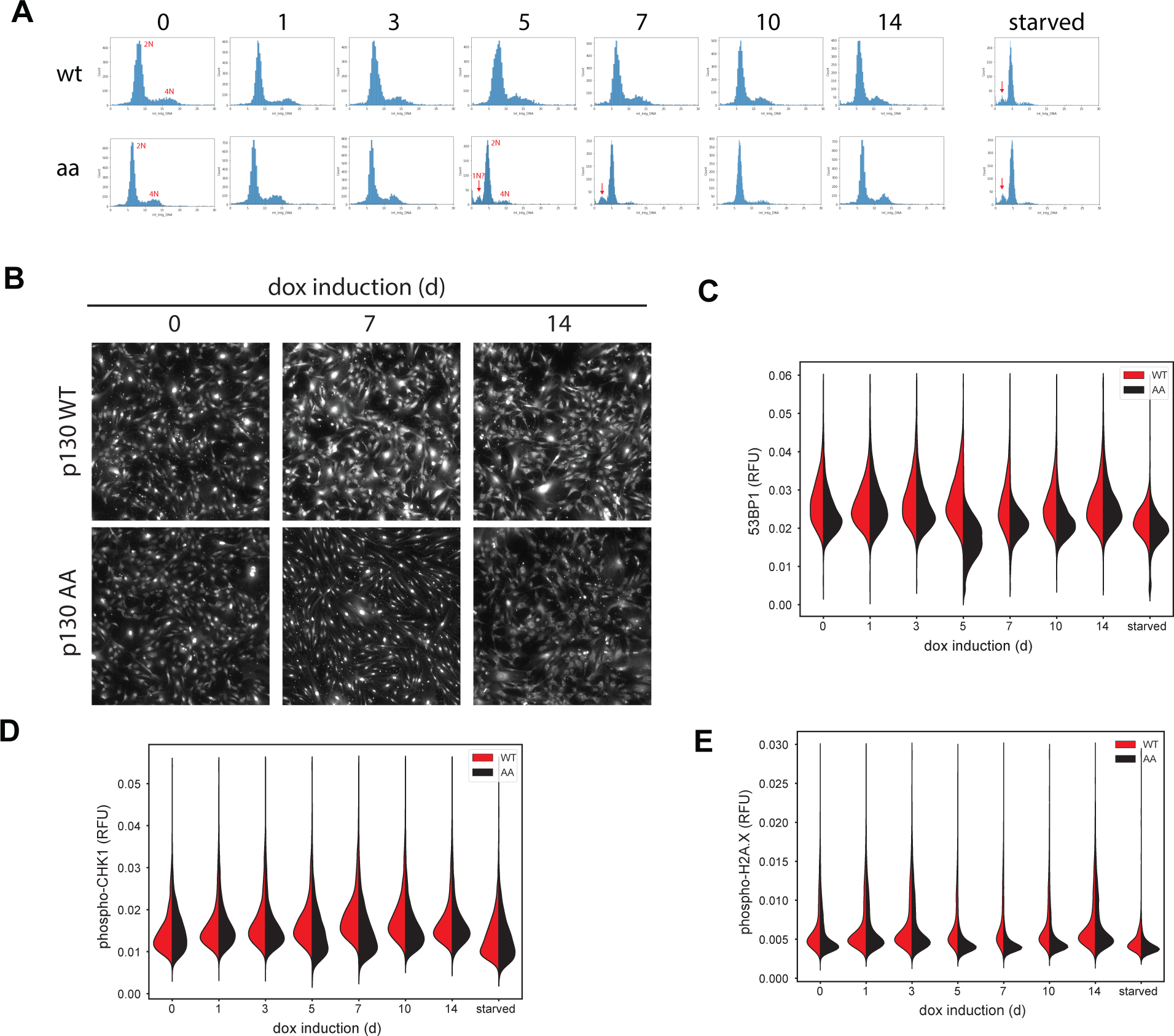
Additional DNA and protein staining in p130(WT) and p130(AA)-expressing cells. ***(A-H)*** Inducible p130 NHF-1 cells were grown in doxycycline for indicated time, then protein expression and protein localization were visualized by 4i staining. ***(A)*** DNA content assessed by DAPI staining. ***(B)*** p130 localization. ***(C)*** total 53BP1 levels***. (D)*** Total phosphor-chk1 levels. ***(E)*** Total phosphor-H2A.X levels.

**Figure S6.**
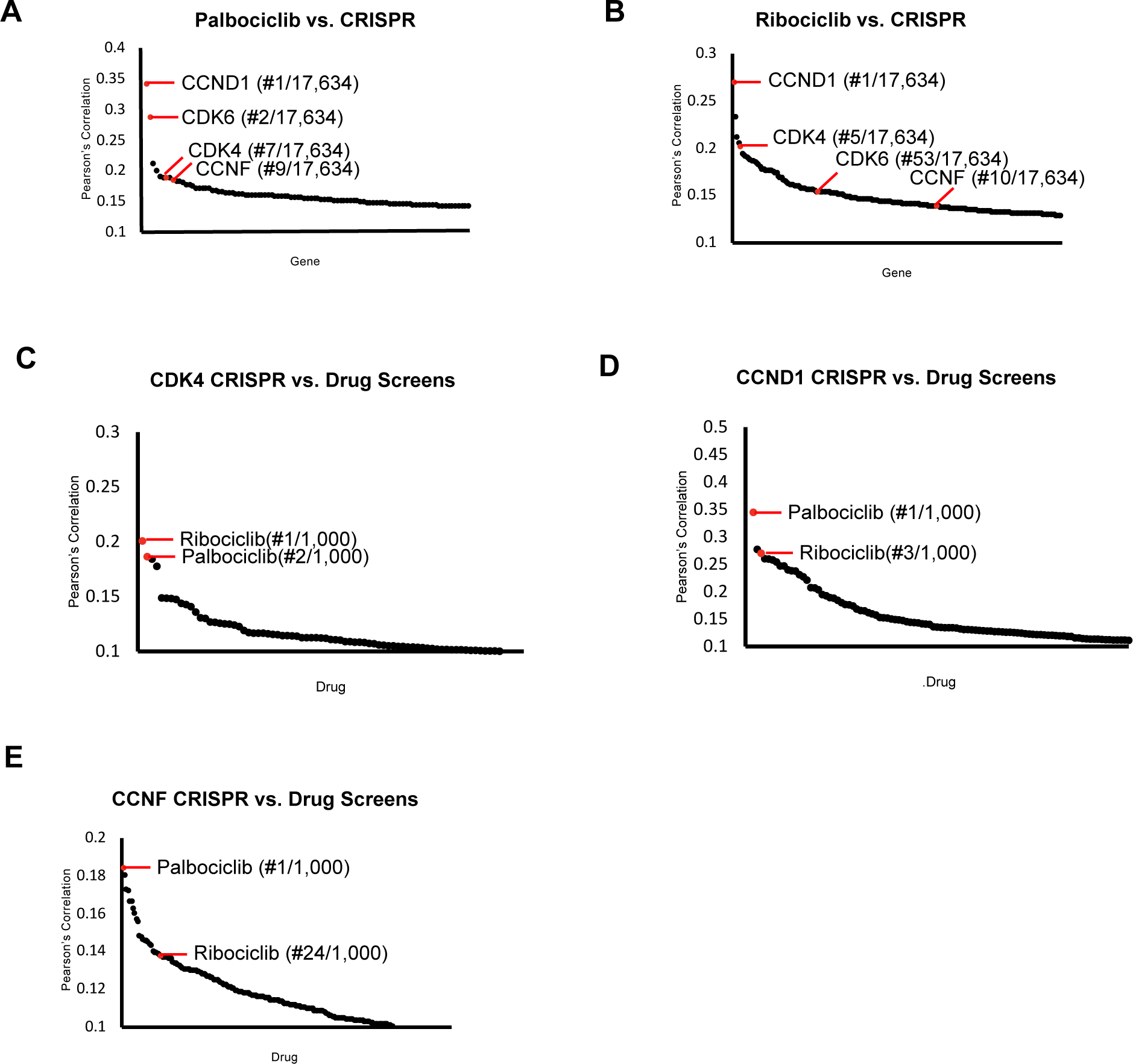
CDK4/6 inhibitors and CCNF knockout correlate highly. Project Achilles Dependency Map datasets were analyzed to determine: ***(A)*** Pearson’s correlation coefficients for gene knockout correlated with Palbociclib treatment. ***(B)*** Pearson’s correlation coefficients for gene knockout correlated with Ribociclib treatment. ***(C)*** Pearson’s correlation coefficients for the correlation between *CDK4* knockout and 1000 drug treatments. ***(D)*** Pearson’s correlation coefficients for the correlation between *CCND1* knockout and 1000 drug treatments. ***(E)*** Pearson’s correlation coefficients for the correlation between *CCNF* knockout and 1000 drug treatments. For (A-E) only the top scoring, most statistically significant associations are shown.

**Table S1.**
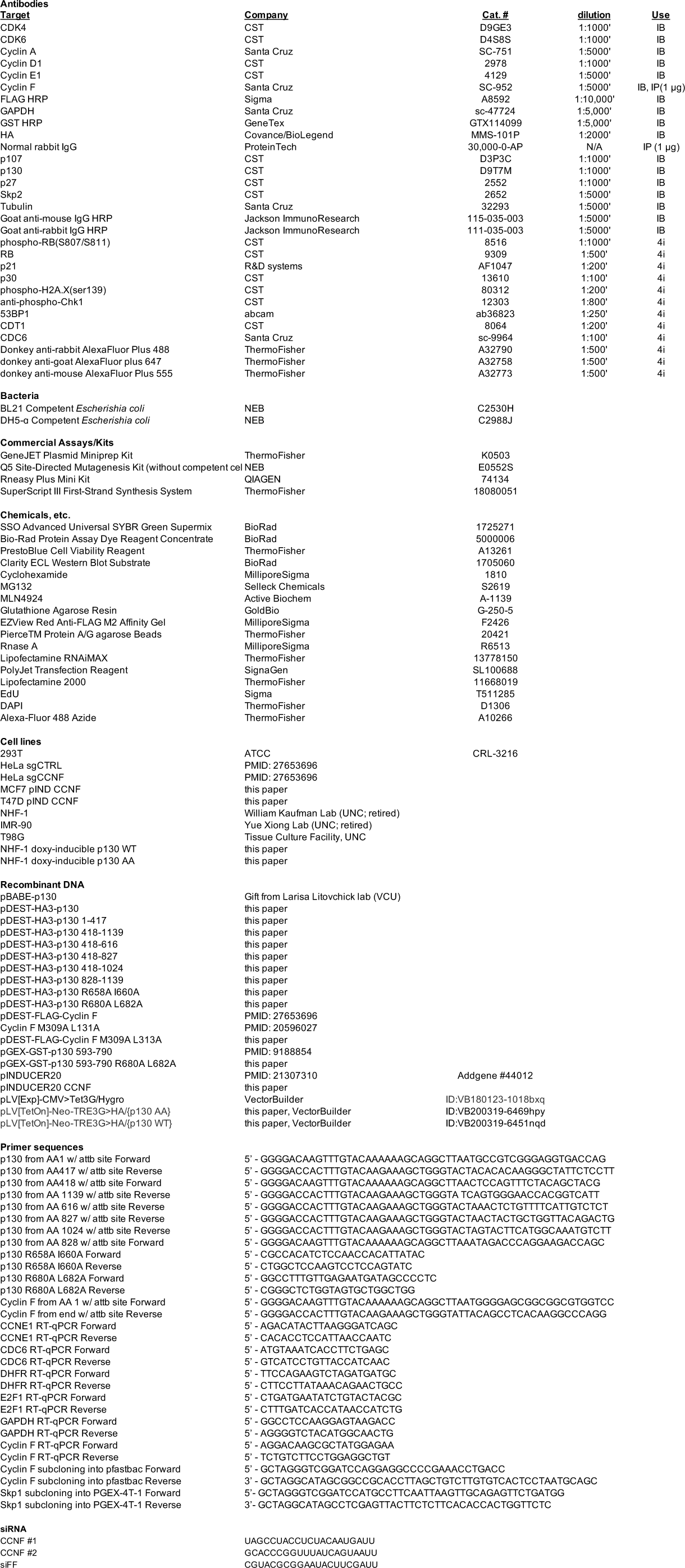
Critical Reagents.

